# Epithelial miR-149-5p up-regulation is associated with immune evasion in progressive bronchial premalignant lesions

**DOI:** 10.1101/2025.02.03.636307

**Authors:** B Ning, DJ Chiu, RM Pfefferkorn, Y Kefella, E Kane, V Reyes-Ortiz, G Liu, S Zhang, H Liu, L Sultan, E Green, M Constant, AE Spira, JD Campbell, ME Reid, X Varelas, EJ Burks, ME Lenburg, SA Mazzilli, JE Beane

**Affiliations:** Department of Medicine, Boston University Chobanian & Avedisian School of Medicine, Boston, MA, USA; Department of Pathology and Laboratory Medicine, Boston University Chobanian & Avedisian School of Medicine, Boston, MA, USA; Department of Medicine, Roswell Park Comprehensive Cancer Center, Buffalo, NY, USA; Department of Biochemistry, Boston University School of Medicine, Boston, MA, USA

## Abstract

The molecular drivers bronchial premalignant lesion progression to invasive lung squamous cell carcinoma are not well defined. Prior work profiling longitudinally collected bronchial premalignant lesion biopsies by RNA sequencing defined a proliferative subtype, enriched with bronchial dysplasia. We found that a gene co-expression module associated with interferon gamma signaling and antigen processing/presentation was down-regulated in progressive/persistent versus regressive lesions within the proliferative subtype, suggesting a functional impact of these genes on immune evasion. RNA from these same premalignant lesions was profiled by microRNA (miRNA) sequencing and a miRNA-gene network analysis identified hsa-miR-149-5p as a potential regulator of this antigen presentation gene co-expression module associated with lesion progression. hsa-miR-149-5p was found to be predominantly expressed in the epithelium and up-regulated in progressive/persistent versus regressive proliferative lesions while targets of this miRNA, the transcriptional coactivator of MHC-I gene expression, *NLRC5*, and the genes it regulates were down-regulated. MicroRNA in situ hybridization of hsa-miR-149-5p in tissue from adjacent fixed biopsies showed that hsa-miR-149-5p was increased in areas of bronchial dysplasia in progressive/persistent versus regressive lesions. Imaging mass cytometry showed that NLRC5 protein expression was decreased in progressive/persistent versus regressive lesions within areas of hyperplasia, metaplasia, and dysplasia. Additionally, basal cells with high versus low levels of NLRC5 were found to be in close spatial proximity to CD8 T cells, suggesting that these cells exhibit increased functional MHC-I gene expression in lesions with low hsa-miR-149-5p expression. Collectively, our data suggests a functional role for hsa-miR-149-5p in bronchial premalignant lesions and may serve as a therapeutic target for PML immunomodulation.

**STATEMENT OF SIGNIFICANCE:** Integrative analysis across bronchial premalignant lesions has identified and localized a potential regulator of immune evasion in progressive/persistent lesions that could be a novel therapeutic target.

## INTRODUCTION

Lung squamous cell carcinoma (LUSC) arises in the bronchial airway and makes up approximately 25% of all lung cancers, the leading cause of cancer-related deaths in the United States and worldwide^1,2^. LUSC is thought to progress from bronchial premalignant lesions (PMLs) that are characterized by aberrant epithelial architecture and abnormal expansion of airway basal cells. The pathologic grading of PMLs has been well documented^3^ and describes progressive epithelial changes including hyperplasia, metaplasia, dysplasia (mild, moderate, and severe), and carcinoma in situ (CIS). As the normal bronchial epithelium is damaged, the differentiated pseudostratified columnar epithelium becomes hyperplastic with the proliferation of basal like cells, or metaplastic with the expansion of squamous cells, and worsens to dysplasia and carcinoma in situ with increased cytologic aberrations including nuclear pleomorphism, increased mitotic figures, and altered cellular differentiation^3–6^. PMLs that progress from normal, hyperplasia or metaplasia to a high-grade PML (moderate/severe dysplasia or CIS) or high-grade PMLs that persists over time are associated with a higher risk for lung cancer development at the site of the PML or elsewhere in the lung^4,7,8^. Only a small fraction of high-grade PMLs, however, progress to CIS or invasive cancer and more than half regress to a lower histologic grade spontaneously without any intervention^4,5^. DNA and methylation profiling of CIS epithelium revealed increased chromosomal instability and somatic mutations in lesions that progress to LUSC, but that spontaneous regression of CIS can occur in cases with known cancer driver mutations^9^. These results suggest that alterations in immune surveillance may be a key determinant of progression to invasive carcinoma.

To gain insight into progression mechanisms, our group and others have profiled the transcriptomes of PMLs obtained from patients at high-risk for developing lung cancer and observed that PML progression is associated with suppression of immune surveillance via multiple mechanisms including impairment of antigen processing and presentation and interferon gamma signaling, increased expression of immune checkpoints and cytokines inhibiting T cell activation, and by promoting the recruitment of myeloid derived tumor suppressor cells^6,10–12^. Our studies have identified a transcriptionally distinct group of PMLs (“proliferative subtype”), based on nine gene co-expression modules, that express high levels of cell cycle and proliferation genes, are enriched with bronchial dysplasia, and have high basal cell and low ciliated cell composition. Persistent or progressive lesions transcriptionally classified as proliferative subtype PMLs have reduced numbers of CD8 T cells and decreased expression of a co-expression module involved in interferon signaling and antigen processing and presentation, termed the antigen presentation gene module, compared with regressive PMLs. Despite the multiple reports of immune evasion in progressive high-grade lesions, little is known about what drives immune-related gene regulation.

In the absence of T cell exhaustion in our cohort, we hypothesized that microRNAs (miRNAs) may be regulating the expression of genes in the antigen presentation gene module associated with PML progression. miRNAs are short non-coding RNAs that suppress gene expression and facilitate transcript degradation through complementary base-pair binding to the 3’ untranslated region of gene transcripts^13^. One study profiling 14 miRNAs observed hsa-miR-34c to be associated with PML histologic changes^14^; however, miRNA expression alterations contributing to bronchial PML progression have not been reported. To address this question, we used sample matched mRNA and miRNA sequencing data from longitudinally collected PML biopsies to probe if miRNA-mediated gene regulation leads to PML progression associated with an immunosuppressive microenvironment. We identified hsa-miR-149-5p as a potential regulator of the antigen presentation gene module associated with proliferative subtype lesion progression. We found hsa-miR-149-5p to be highly expressed in the epithelium, up-regulated in progressing/persistent compared to regressing proliferative subtype PMLs, and a regulator of major histocompatibility complex class I (MHC-I) and antigen processing genes via *NLRC5* that influences recruitment of CD8 T cells. Our results suggest that hsa-miR-149-5p may be a potential therapeutic target for preventing PML progression to lung cancer.

## RESULTS

### miRNA-gene network analysis identifies miRNAs associated with gene co-expression modules that define PML molecular subtypes

As miRNAs have been shown to be important in carcinogenesis^15,16^, we sought to construct a miRNA-gene network from bronchial PMLs to elucidate how miRNAs may be negatively regulating target genes associated with PML progression. To construct a miRNA-gene network, we generated high quality miRNA sequencing data from n=156 endobronchial biopsies from 28 patients that had matching bulk RNA sequencing data^10^ **(Figure 1A)**. This subset was representative of the larger previously published cohort^10^, based on the observed associations between lesion molecular subtype, smoking status, and histology (chi-square test p-value < 0.05) (**Table 1 and Supplementary Table 1)**. The paired miRNA and mRNA expression data was used to construct the network where nodes are miRNAs and genes and edges represent significant negative correlations between miRNAs and their predicted target genes. After removing miRNA/gene nodes without any connections, the miRNA-gene network consisted of 14744 edges between 464 miRNAs and 2296 genes. The median number of connected target genes for each miRNA was 18. Next, we filtered edges **(Figure 1B**; see **Methods**) to identify module associated miRNAs that may have central gene-expression regulatory roles within the miRNA-gene network in each of the nine previously described^10^ gene modules that define the PML molecular subtypes (**Supplemental Table 2**). For each gene module, we identified and retained only the connections from miRNAs whose associations with the module target genes are stronger (more negative) than with non-module target genes and where the significant negatively correlated target genes are significantly enriched within the module. After excluding nodes without any connections and summarizing miRNA-gene connections to the gene module level, we obtained a bipartite directed network consisting of 321 miRNA nodes targeting genes in the nine gene modules (**Figure 2A**; **Supplementary Table 3**). The connectivity of the miRNA-gene module network did not show a dependence on the number of genes in a module and 94 miRNAs had connections to more than one gene module.

**Figure 1.**
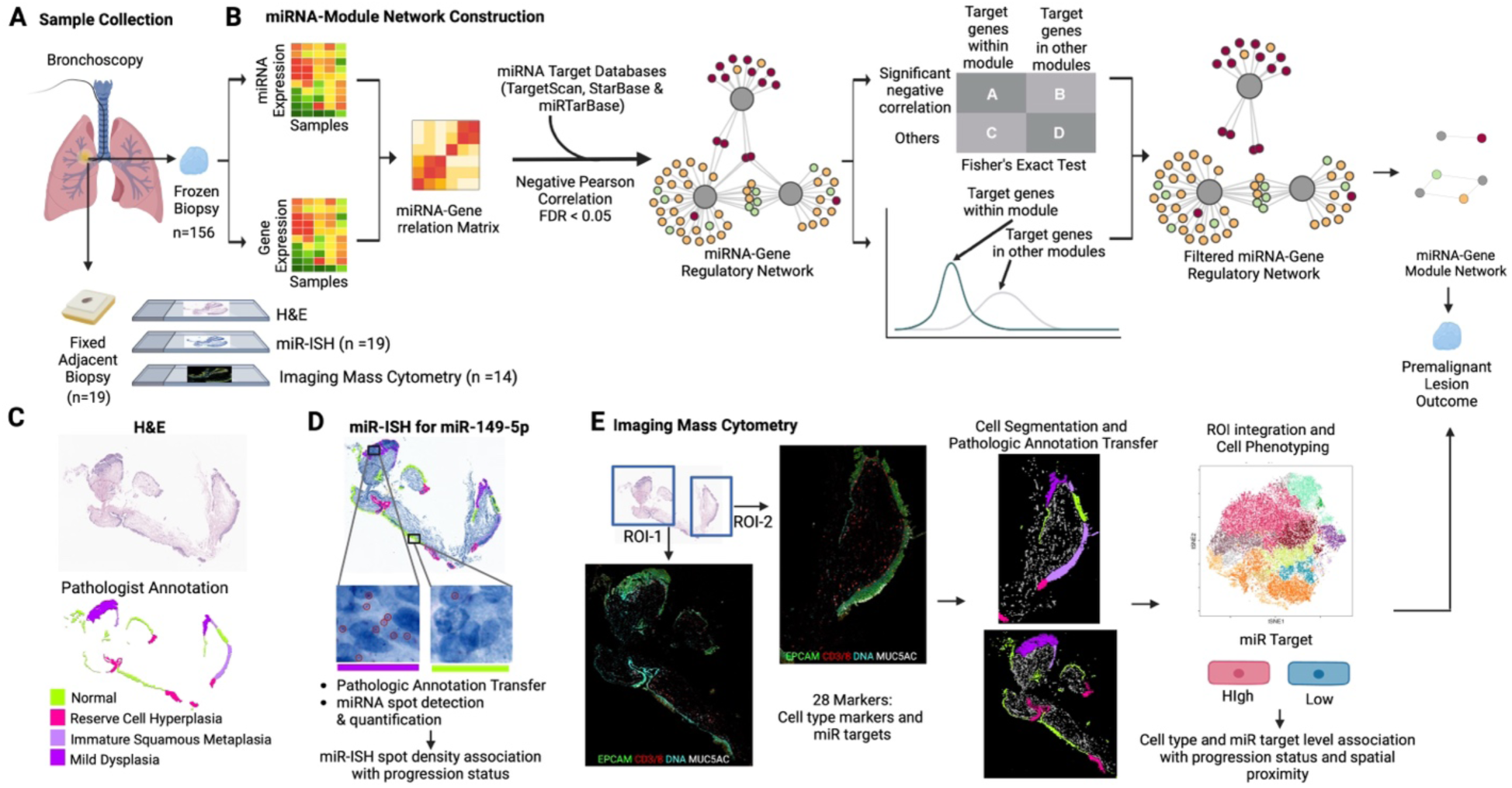
Study overview. (**A**) RNA isolated from 156 frozen endobronchial biopsies (n=28 patients) was profiled by miRNA sequencing. These samples also had paired mRNA sequencing data. Among the 156 biopsies, 19 adjacent FFPE biopsies, transcriptionally defined as the proliferative molecular subtype, were sectioned and stained to assess spatial localization of key findings. (**B**) The paired miRNA-mRNA sequencing data was used to construct a miRNA-gene network based on database evidence of miRNA target genes and significant negative correlation between miRNA and gene expression patterns (gray circles are miRNAs and red, green, and yellow circles are genes where the color represents their membership in previously described gene co-expression modules that define PML molecular subtype). Target information was obtained by combining a sequence-based prediction database and experimentally validated databases. Pearson correlation was calculated from expression residuals for each miRNA-mRNA pair, and those with significant negative Pearson correlation coefficients (FDR <= 0.05) were selected. Next, we filtered edges to identify miRNAs that may have central gene expression regulatory roles within the miRNA-gene network in each gene module. For each gene module, we identified and retained only the connections from miRNAs where 1) there are more significantly negative correlated target genes in the module than in other modules (Fisher-exact test, odds ratio>1) and 2) the Fisher-Z transformed Pearson correlation coefficient densities with its targets within the module were more negative than its targets in other modules (t-test, t < 0). miRNAs with significant results from both tests (FDR <= 0.05) were considered module associated miRNAs and other miRNAs were removed from the network. (**C**, **D** and **E**) For each FFPE biopsy, three consecutive sections were stained for different purposes: (**C**) H&E to annotate the histologic grades present in the epithelial tissue, (**D**) chromogenic miRNA in situ hybridization (miRNA-ISH) to quantify hsa-miR-149-5p expression and localization, and (**E**) imaging mass cytometry (IMC) using 28 metal-tagged antibodies to identify epithelial and immune cell types and hsa-miR-149-5p target protein expression and localization. Single or multiple regions of interest (ROI) per sample were selected for IMC ablation in 14 samples. The epithelial histology in both miRNA-ISH and IMC images were annotated based on the corresponding adjacent H&E section. Created in BioRender. (2025) https://BioRender.com/g78j632

**Figure 2.**
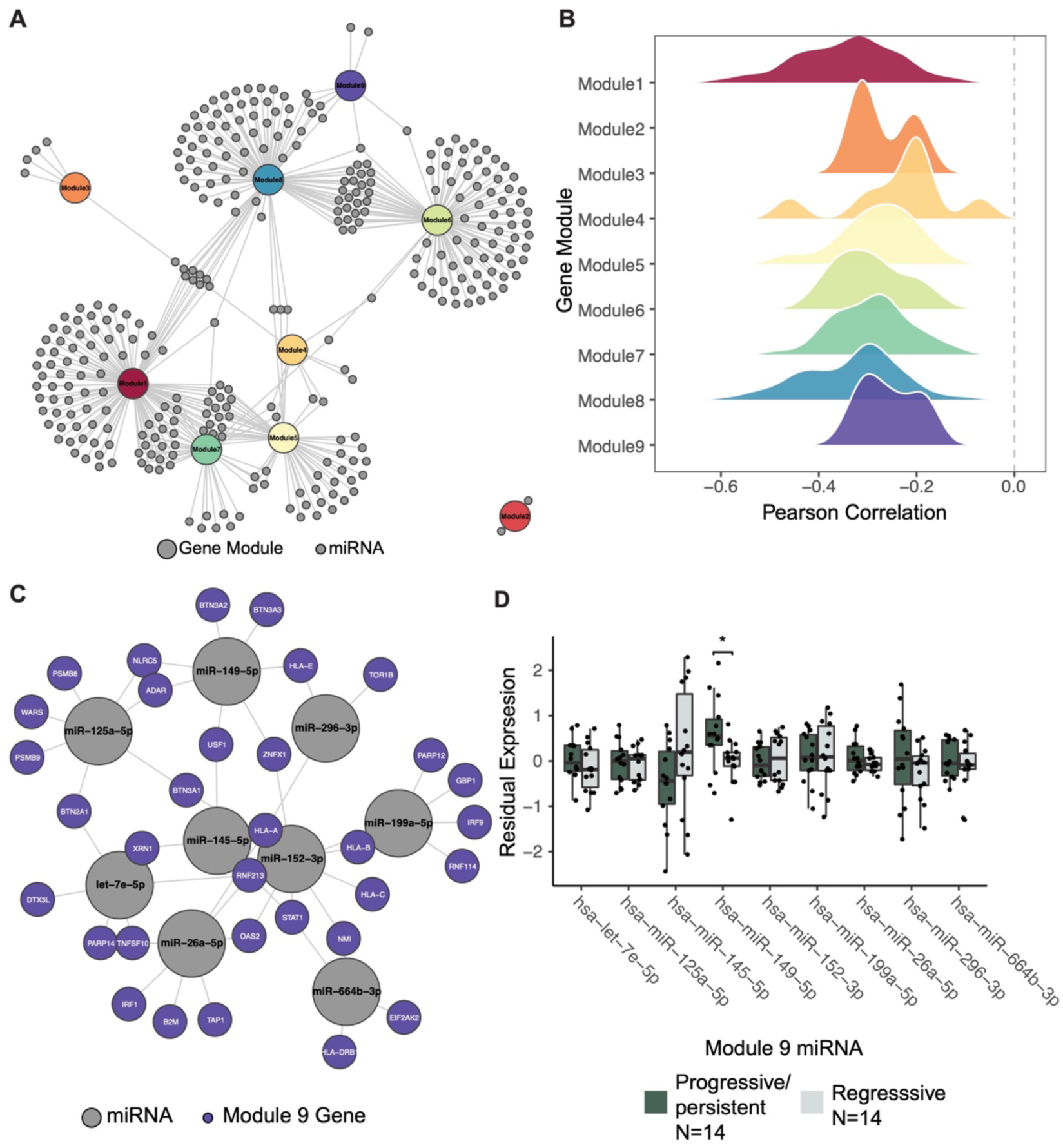
miRNA-gene network analysis identifies miRNAs associated with the antigen presentation gene module down-regulated in progressive proliferative subtype PMLs. (**A**) miRNA-gene module network plot shows 321 miRNAs connected to 9 gene modules where grey circles are miRNAs and colored circles are the gene modules. Edges reflect a miRNA may have strong suppressive regulatory role over the predicted target genes within the connected gene module based on the two statistical test filters. (**B**) Pearson correlation densities between the module associated miRNAs connected to each gene module in the miRNA-gene module network and the corresponding gene module GSVA score. (**C**) miRNA-gene network plot of miRNAs connected to the antigen presentation gene module (Module 9, purple circle in (**A**)). Edges show the significant negative correlations between the miRNAs (grey circles) and the predicted target genes (purple circles). (**D**) Boxplots of residual expression levels of the 9 miRNAs specifically connected to module 9 in proliferative subtype biopsies stratified by their outcome status (progressive/persistent samples in dark green and regressive samples in light green). *P < 0.05, determined by linear mixed effect model.

**Table 1.** Clinical annotations for endobronchial biopsies that were profiled by miRNA sequencing stratified by PML molecular subtypes. Statistical tests for categorical clinical variables (dysplasia grade, smoking status, progression status and batch) were conducted using Chi-square tests. Statistical tests for continuous variables (TIN) were compared using two-sided Student’s t-tests. Percentages are reported for categorical variables and mean/standard deviations are reported for the continuous variable.

|  | **Proliferative**<br>**(n=46)** | **Inflammatory**<br>**(n=27)** | **Secretory**<br>**(n=46)** | **Normal**<br>**(n=37)** |  |
| --- | --- | --- | --- | --- | --- |
| **Dysplasia Grade** |  |  |  |  | **chi=38.50,**<br>**p=0.0033** |
| **Normal** | 3 (6.5) | 4 (14.8) | 12 (26.1) | 10 (27.0) |  |
| **Hyperplasia** | 4 (8.7) | 8 (29.6) | 7 (15.2) | 6 (16.2) |  |
| **Metaplasia** | 8 (17.4) | 8 (39.6) | 9 (19.6) | 12 (32.4) |  |
| **Mild Dysplasia** | 5 (10.9) | 1 (3.7) | 8 (17.4) | 2 (5.4) |  |
| **Moderate Dysplasia** | 19 (41.3) | 4 (14.8) | 7 (15.2) | 5 (13.5) |  |
| **Severe Dysplasia** | 7 (15.2) | 1 (3.7) | 1 (2.2) | 1 (2.7) |  |
| **UNK** | 0 (0.0) | 1 (3.7) | 2 (4.4) | 1 (2.7) |  |
| **Smoking Status (genomic prediction)** | 40 (87.0) | 11 (40.7) | 35 (76.1) | 12 (32.4) | **chi=35.20,**<br>**p < 0.001** |
| **Progression Status** |  |  |  |  | **chi=24.37,**<br>**p=0.0038** |
| **Progression/persistent** | 14 (30.4) | 4 (14.8) | 14 (30.4) | 5 (13.6) |  |
| **Regressing** | 14 (30.4) | 3 (11.1) | 5 (10.9) | 3 (8.1) |  |
| **Normal/Stable** | 7 (15.2) | 12 (44.4) | 8 (17.4) | 14 (37.8) |  |
| **UNK** | 11 (23.9) | 8 (29.6) | 19 (41.3) | 15 (40.5) |  |

To validate that our network captures biologically relevant miRNA-gene relationships, for each module, we confirmed that the correlation density between the expression levels of module associated miRNAs and a module metagene score computed via GSVA were left-shifted from zero (**Figure 2B**). We also observed a significant negative correlation pattern between the GSVA score of a gene module and the GSVA score calculated from the module associated miRNAs (i.e. negative correlation along the diagonal line in **Extended Data Figure 1**; FDR < 0.05, Pearson correlation), except for Module 2. Moreover, the connections between miRNA and gene modules in the network reflect multifaceted functions of miRNAs demonstrated by previous studies. ^17,18^ For example, miR-200a/b-3p are connected to gene modules with genes associated with the extracellular matrix and the immune activation^19,20^ (module 1 and 8**; Supplementary Table 3**). These results suggest that our miRNA-gene module network reveals biologically meaningful regulatory relationships between miRNAs and the predicted target genes.

### Genes in the PML progression-associated antigen presentation module are connected to hsa-miR-149-5p

The antigen presentation gene module (module 9), enriched with genes associated with interferon signaling and antigen presentation and processing, has previously^10^ been shown to be down-regulated among proliferative subtype progressive/persistent compared to the regressive PMLs. To identify miRNAs that potentially contribute to the PML progressive phenotype, we used the miRNA-gene module network to identify 9 miRNAs (hsa-let-7e-5p, hsa-miR-125a-5p, hsa-miR-145-5p, hsa-miR-149-5p, hsa-miR-152-3p, hsa-miR-199a-5p, hsa-miR-26a-5p, hsa-miR-296-3p and hsa-miR-664b-3p) that were significantly negatively correlated with 33 out of the 112 module 9 genes (**Figure 2C**). For each of these 9 miRNAs, the miRNA-gene correlations with predicted target genes in module 9 were significantly more negative than correlations with predicted target genes in the other gene modules, and the significant negatively correlated target genes were enriched within module 9 (FDR < 0.05, t-test and Fisher-exact test; **Supplementary Table 4**). Next, we examined whether the expression levels of these miRNAs are associated with the progressive status of proliferative subtype PMLs. hsa-miR-149-5p was the only significantly up-regulated miRNA in progressive/persistent (n=14) versus regressive (n=14) proliferative subtype PMLs (p-value < 0.05, linear mixed-effect model; **Figure 2D and Supplementary Table 5**), suggesting its higher expression is associated with worse lesion prognosis in concordance with its previously reported oncogenic role in different cancer settings^21^.

### hsa-miR-149-5p targets MHC-I genes by suppressing NLRC5 expression

Next, we sought to elucidate how hsa-miR-149-5p up-regulation contributes to lesion progression by investigating the 7 genes in module 9 connected to the miRNA in the network analysis (**Figure 2C**). We compared the expression levels of these genes between the proliferative subtype progressive/persistent (n=15) versus the regressive (n=15) PMLs from the discovery cohort in our prior work^10^ (GSE109473). All seven targets were down-regulated in the progressive/persistent PML samples, and the expression level of three target genes, *BTN3A3*, *HLA-E*, and *NLRC5*, are significantly lower in the progressive/persistent proliferative PMLs than in the regressive ones (p-value < 0.05, linear mixed-effect model), (**Figure 3A and Supplementary Table 6**). The lower expression of these genes among PMLs that progress is also observed in the proliferative PML samples from our validation cohort (n=7 progressive/persistent and n=12 regressive) and in an independent dataset from Merrick *et. al*. (n=32 progressive/persistent and n=15 regressive, GSE114489) (**Supplementary Table 6**).

**Figure 3.**
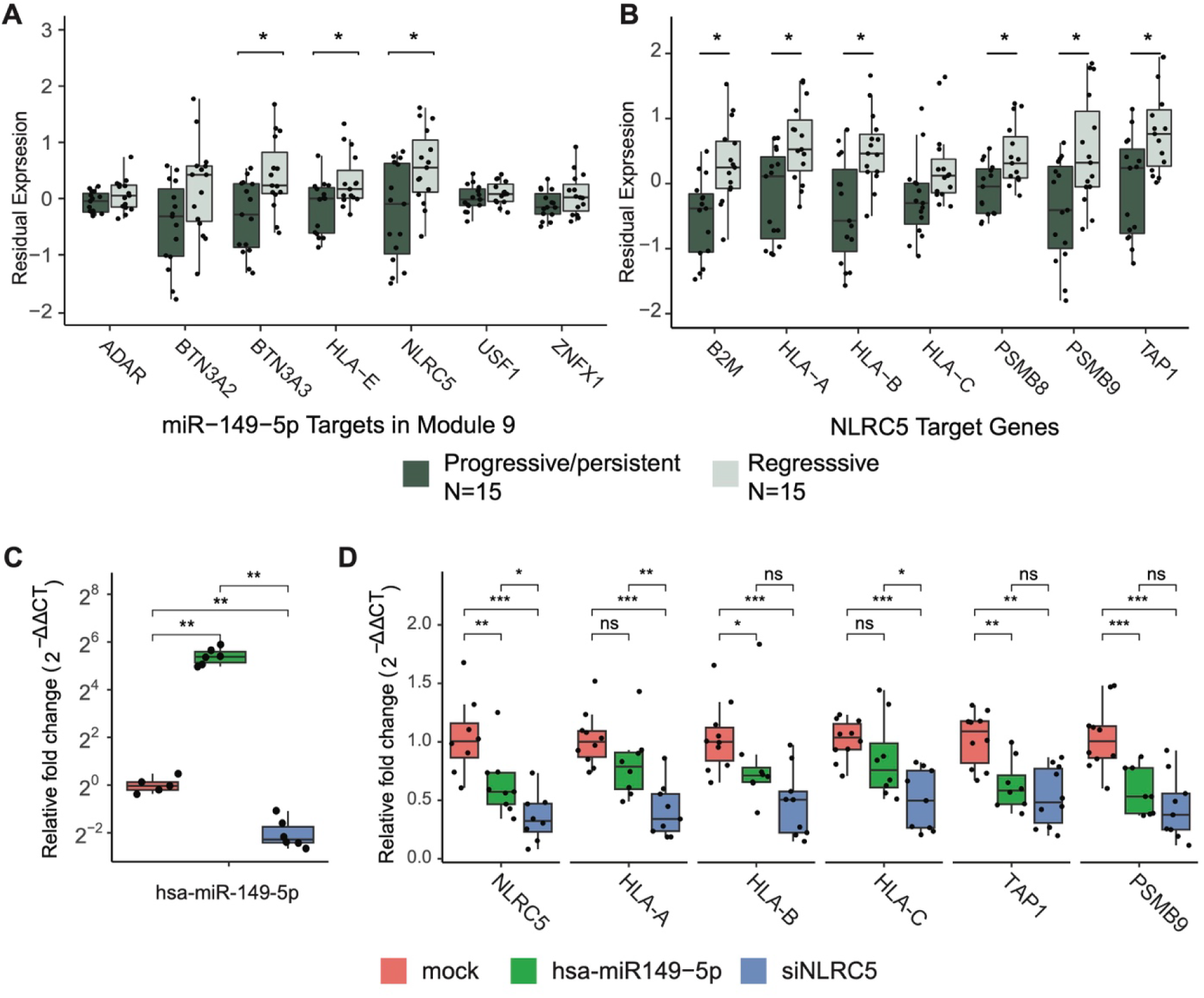
Hsa-miR-149-5p and NLRC5 target genes are associated with PML progression and change when hsa-miR-149-5p and NLRC5 are modulated in vitro. Boxplots of residual expression levels of between progressive/persistent (dark green) and regressive (light green) Proliferative subtype PMLs from the discovery cohort (GSE109473) for (**A**) hsa-miR-149-5p significant negatively correlated target genes in module 9 and (**B**) NLRC5 target genes. SW900 cells were transfected with mock control (red orange), hsa-miR-149-5p (green) or siNLRC5 (blue) for 48 hours. Relative gene expression of (**C**) miR-149 and (**D**) NLRC5 and its downstream targets (HLA-A, HLA-B, HLA-C, TAP1, and PSMB9) were detected by qRT-PCR (6 to 10 replicates per condition). PPIA was used as the reference gene to calculate relative fold change values. Data indicate median with IQR, and whiskers indicate minimum and maximum measurement. P values were determined using mixed effect linear models for (**A**) and (**B**); and two-sided Wilcoxon tests for (**C**) and (**D**). *P <= 0.05, **P <= 0.01, ***P <= 0.001; ns, no significance.

To further validate the association between hsa-miR-149-5p and module 9, we leveraged three airway and lung tumor datasets with sample-matched miRNA and gene expression profiles: airway brushing samples (n=87) obtained during the same bronchoscopy procedure as the endobronchial biopsies used to derive the miRNA-RNA network cohort, TCGA LUSC resection samples^22^ (n=475), and airway brushings from the Airway Epithelial Gene Expression in the Diagnosis of Lung Cancer (AEGIS) trials^23,24^ (n=341). Module 9 genes were significantly enriched among genes that were negatively correlated with hsa-miR-149-5p in all datasets (p-value < 0.001, GSEA; **Extended Data Figure 2A**). Also, 47.7% of the genes in the leading edge driving the signal were overlapping across four studies, including four of the hsa-miR-149-5p predicted target genes in module 9 (*BTN3A2, BTN3A3, HLA-E* and *NLRC5*; **Extended Data Figure 2B**). These observations suggest hsa-miR-149-5p may promote the progressive PML phenotype by specifically suppressing target genes in the antigen presentation gene module.

Among the hsa-miR-149-5p target genes significantly down-regulated in progressive/persistent lesions, we found *NLRC5* to be particularly interesting because previous studies showed that NLRC5 is a member of the NOD-like receptor (NLR) family and is a transcriptional activator of the MHC-I genes^25^. *NLRC5* is a predicted target for hsa-miR-149-5p in two out of the three miRNA target databases^26–28^ used in our network construction, and was negatively correlated to hsa-miR-149-5p in the three airway and lung tissue datasets (p <= 0.05, Pearson correlation; **Extended Data Figure 3A-C**). We previously reported that depletion of CD8 T cells is associated with PML progression^10^, and hypothesized this could partially be due to hsa-miR-149-5p repressing the expression of the MHC-I genes by down-regulating *NLRC5* expression. Seven genes (*B2M, HLA-A, HLA-B, HLA-C, PSMB8, PSMB9*, and *TAP1*) were identified to be directly regulated by NLRC5 based on literature reviews^29,30^. NLRC5 promoter-binding at the mouse orthologues of these genes can be observed via ChIP-seq experiments from mice T cells^31^. The 7 *NLRC5* target genes are also within module 9, and the expression levels for all, except *HLA-C*, were significantly down-regulated within the proliferative subtype progressive/persistent versus regressive PMLs in the previously published discovery cohort (**Figure 3B and Supplementary Table 7**). We further validated the downregulation of *NLRC5* target genes in progressive/persistent versus regressive PMLs in the proliferative subtype PMLs from our validation cohort and in the samples from Merrick *et. al.* (GSE114489) (**Supplementary Table 7**).

qPCR experiments in SW900, lung squamous cancer cell lines, transfected with hsa-miR-149-5p, siNLRC5, or a mock sequence, demonstrated that transfection with hsa-miR-149-5p significantly increased hsa-miR-149-5p and decreased *NLRC5* and its target genes, *HLA-B*, *TAP1*, and *PSMB9*, compared to mock. Transfection with siNLRC5 significantly decreased *NLRC5* and its target genes *HLA-A/B/C*, *PSMB9*, and *TAP1* (**Figure 3C and 3D**; two-sided Wilcoxon test p-value < 0.05). The experiments also suggest a potential negative feedback loop between *NLRC5* and hsa-miR-149-5p as *NLRC5* was decreased more strongly by experiments with siNLRC5 than hsa-miR-149-5p alone and siNLRC5 significantly decreased hsa-miR-149-5p expression compared to mock (**Figure 3C**; two-sided Wilcoxon test p-value < 0.01). The computational and qPCR results suggest that hsa-miR-149-5p may promote lesion progression by suppressing *NLRC5* and MHC-I gene expression.

### hsa-miR-149-5p expression is enriched within epithelial cells

Interpreting the potential role of hsa-miR-149-5p in lesion progression is dependent on the cell type specificity of its expression and its targets. To identify the cell type specificity of hsa-miR-149-5p expression, we examined the correlation pattern between hsa-miR-149-5p and canonical cell type markers within our lesion biopsy samples. We observed significant positive correlations between hsa-miR-149-5p and the basal cell marker, *KRT5*^32^ (FDR < 0.001, Pearson correlation) and the peri-goblet cell marker, *CEACAM5*^33^ (FDR < 0.01, Pearson correlation; **Figure 4A**). In comparison, hsa-miR-34b-5p and hsa-miR-449c-5p, which are expressed highly in mucociliary epithelia, were positively correlated with *FOXJ1*, and hematopoiesis essential hsa-miR-150-5p was positively correlated with B and T cell markers^34^ (**Figure 4A)**. We also utilized cell type-specific transcriptomic data from the FANTOM5 project^35^ and ranked samples based on the normalized expression level of hsa-miR-149-5p and identified whether samples derived from epithelial or lymphoid cell types were enriched among samples with high hsa-miR-149-5p expression. Samples from epithelial cells (n=87) were positively enriched among samples with high hsa-miR-149-5p expression (p < 0.001, GSEA), while samples from immune cells (n=37) were negatively enriched (p < 0.01, GSEA; **Figure 4B**). Additionally, the expression of hsa-miR-149-5p and *NLRC5* in the FANTOM5 data were significantly negatively correlated between samples from epithelial cells (p < 0.01, Pearson Correlation; **Figure 4C**), but not in samples from immune cells (**Extended Data Figure 4**), suggesting that that the suppression of *NLRC5* by hsa-miR-149-5p may be specific to epithelial cells. The highly enriched expression of hsa-miR-149-5p and negative regulation relationship between hsa-miR-149-5p and *NLRC5* within the epithelial cells suggested that hsa-miR-149-5p may be involved in epithelial cell mediated immune evasion in early lung cancer.

**Figure 4.**
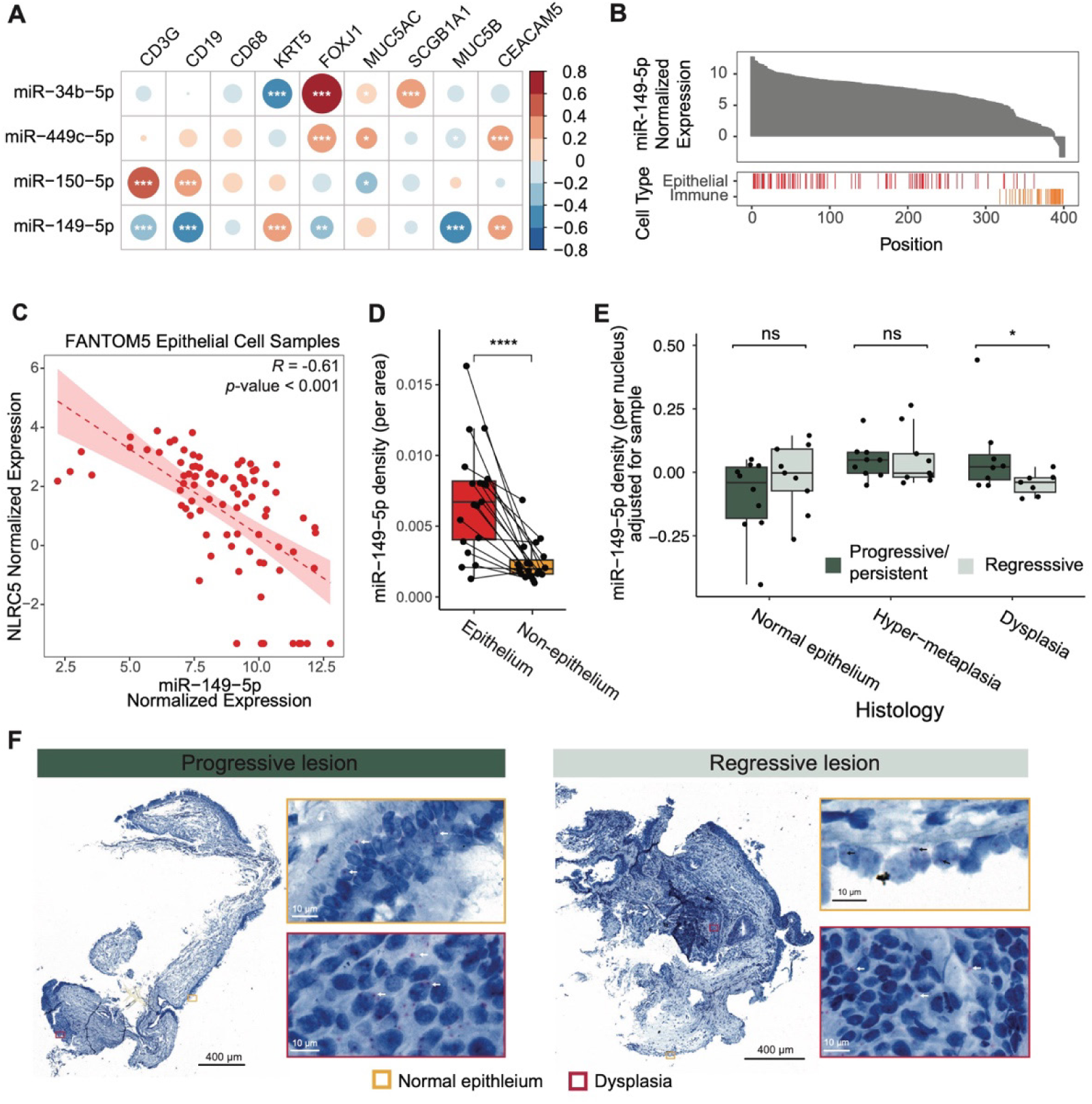
hsa-miR-149-5p is highly expressed within the epithelium and associates with PML progression. **(A)** Bubble plot showing the correlation between the residual expression level of miR-149-5p and cell-type marker genes, including *CD3G* (T cells), *CD19* (B cells), *CD68* (macrophages), *KRT5* (basal cells), *FOXJ1* (ciliated cells), *MUC5AC* (goblet cells), *SCGB1A1* (club cells), *MUC5B* (secretory cells), and *CEACAM5* (peri-goblet cells). Correlation results for hsa-miR-34b-5p, hsa-miR-449c-5p and hsa-miR-150-5p are shown as controls. (**B)** Enrichment of hsa-miR-149-5p across cell-type compartments in the FANTOM5 project. Bar chart (top) shows the normalized expression levels of hsa-miR-149-5p in the FANTOM5 samples (n = 399). Vertical bars (bottom) indicate the position of FANTOM5 samples derived from the epithelial (red) or immune (orange) cell compartments. (**C)** Scatter plot of the Pearson correlation between the normalized expression levels of hsa-miR-149-5p and NLRC5 within the FANTOM5 samples derived from the epithelial cell compartment (n = 87). The dashed red line represents the linear regression fit and the shaded region indicates the 95% confidence interval. (**D-F**) The hsa-miR-149-5p density was detected by miRNA-ISH from proliferative subtype PMLs (n = 19; 10 progressive/persistent, 9 regressive). (**D**) Boxplot showing the hsa-miR-149-5p density per area between the epithelium and non-epithelium regions. (**E**) hsa-miR-149-5p density per nucleus between the progressive/persistent and regressive PMLs within regions of normal epithelium, regions with hyperplasia or metaplasia (hyper-metaplasia), and regions of bronchial dysplasia. (**F**) Representative miRNA-ISH staining of hsa-miR-149-5p for progressive/persistent PMLs (top) and regressive PMLs (bottom). Arrows indicate stained hsa-miR-149-5p within epithelium. Boxplots indicate median with IQR and whiskers indicate minimum and maximum measurement in (**D**) and (**E**). P values were FDR (Benjamini & Hochberg) adjusted and determined by t-test for correlation significance for (**A**), the two-sided paired Wilcoxon test for (**D**), and a linear mixed effect model for (**E**). * P <= 0.05, **P <= 0.01, ***P <= 0.001, ****P <= 0.0001; ns, no significance.

### Spatial localization of hsa-miR-149-5p in PMLs showed higher expression in the epithelium that was associated with PML progression

To localize and validate the computational findings, consecutive tissue sections from proliferative subtype endobronchial biopsies underwent staining with H&E to annotate the histologic grades present in the epithelium and chromogenic miRNA in situ hybridization (miRNA-ISH) with a hsa-miR-149-5p probe to measure hsa-miR-149-5p expression and localization (**Figure 1A**, **Figure 1C-D, Supplemental Table 8**). An expert pathologist annotated the histologic grades of the epithelium in each digitized whole slide H&E image, and these annotations were transferred to the miRNA-ISH and grouped as follows: 1) normal; 2) hyperplasia, immature squamous metaplasia, and squamous metaplasia (denoted as hyper-metaplasia); and 3) squamous dysplasia. The worst histologic grade observed in the H&E matched the clinical diagnosis in 81% of biopsies, suggesting that many features in the clinical specimens were retained in subsequent sections (**Supplemental Table 8, 9**). A pixel classifier was trained to detect hsa-miR-149-5p signal spots in the miRNA-ISH data and its performance was validated on regions (16726 +/- 4332 mm^2^) containing manual spot annotations for each sample. The classifier showed high recall (0.84 +/- 0.06), precision (0.75 +/- 0.14), and F1 (0.78 +/- 0.08) measurement that was not associated with lesion outcome (**Extended Data Figure 5A, 5B**). The detected hsa-miR-149-5p spots per area were higher in the epithelium compared to non-epithelium (p < 0.001, two-sided paired Wilcoxon test) (**Figure 4D**), supporting the computational results showing high hsa-miR-149-5p expression correlation with epithelial markers. We also observed higher hsa-miR-149-5p spot density in epithelial areas with squamous dysplasia in the progressive/persistent compared to regressive PMLs (p=0.05, linear mixed-effect model) (**Figure 4E, 4F**). These results confirm that hsa-miR-149-5p is expressed more highly in epithelial cells and that expression differences observed between progressive/persistent versus regressive PMLs may occur within regions of bronchial dysplasia.

### IMC data shows differential cell composition associated with histology and correlated protein expression between NLRC5 and its downstream targets

The transcriptomic and miRNA-ISH data suggest that hsa-miR-149-5p is more highly expressed in the epithelium, so we sought to confirm that the protein expression between NLRC5 and its downstream targets was correlated within epithelial cells via IMC. Using consecutive tissue sections from those used in the miRNA-ISH experiment, we ablated regions of interest (ROIs) containing epithelial and non-epithelial tissue, and cells from these regions were separated using expression of broad cell type markers (KRT5, E-Cadherin, CD20, CD3, HLA-DR, αSMA, and VIM) coupled with pathologist annotation described above (**Extended Data Figure 6A, 6B)**. The 41,147 high quality cells from the epithelial predominant tissue regions were used to identify 13 cell clusters (**Figure 5A**) containing epithelial and immune cell types based on expression of canonical cell markers (**Figure 5B**). Each cell cluster had cells from at least 13 samples (out of 14 samples tested). We did not, however, detect significant associations between cell cluster abundance and smoking status (FDR>0.05, sccomp differential composition test, **Extended Data Figure 7A, 7B**), potentially due to the underrepresentation of former smokers. The expected location of cell types in the IMC images helped confirm the cluster labels. For example, clusters labeled as basal cells were found close to the basement membrane while clusters labeled as fully differentiated cells (i.e., MUC5AC+ goblet cells), were found at the apical surface in both normal and hyper-metaplastic regions (**Extended Data Figure 9A**). Differential composition analysis showed that fully differentiated cells, including MUC5B+ secretory cells and FOXJ1+ ciliated cells, are decreased in regions of hyper-metaplasia and dysplasia, whereas intermediate state cells including CEACAM5+ peri-goblet cells and CEACAM5+SCGB1A1+ secretory cells, are increased (FDR<0.001, sccomp differential composition test, **Figure 5C, Extended Data Figure 7C**). The infiltration of immune cells, including CD8 T cells, CD4 T cells, NK cells, are significantly increased in dysplastic regions (FDR<0.001, sccomp differential composition test, **Figure 5C, Extended Data Figure 7C**).

**Figure 5.**
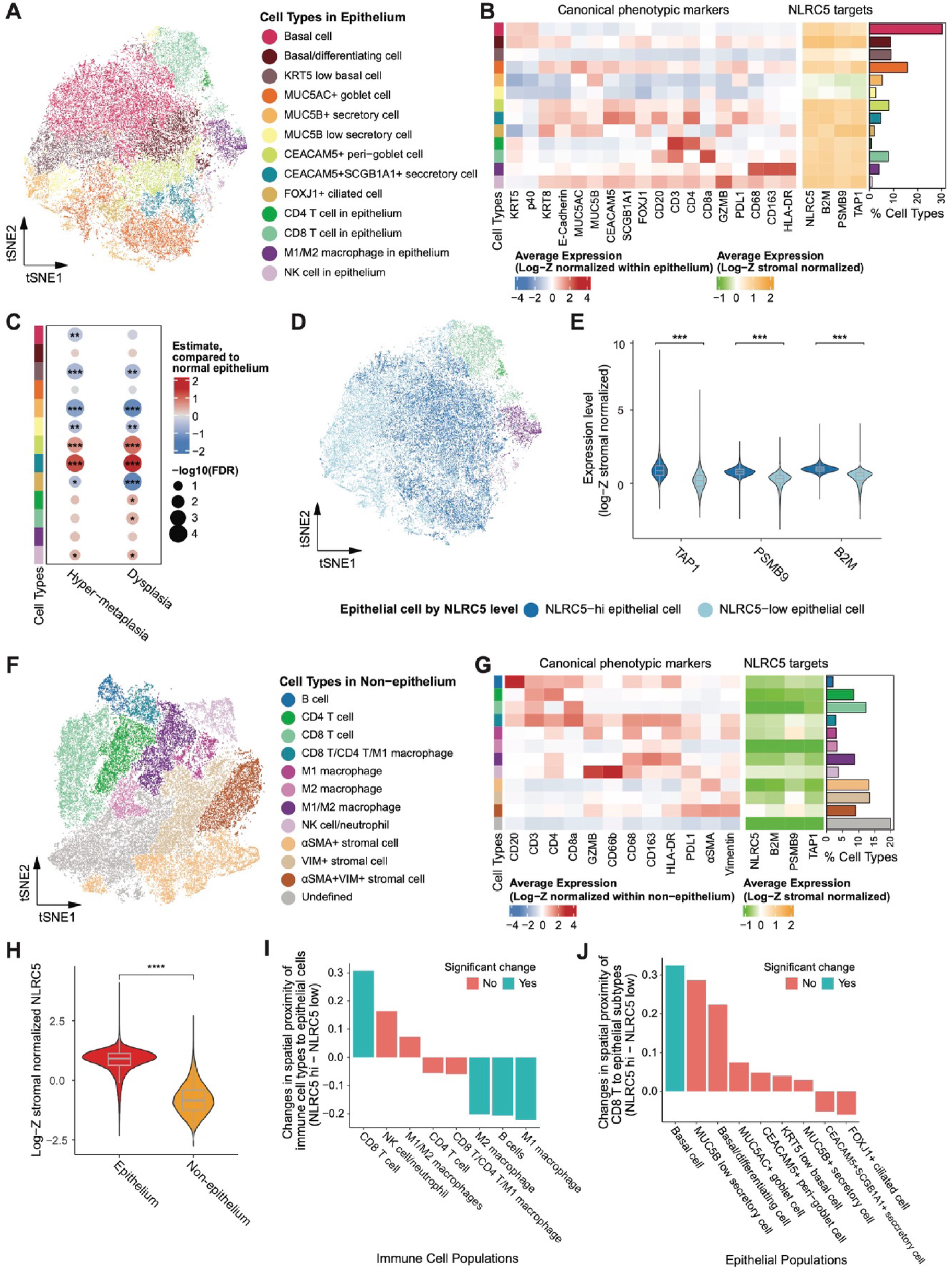
Protein expression of NLRC5 and its targets are correlated and basal cells with increased NLRC5 expression are in close spatial proximity to CD8 T cells. (**A**) t-distributed Stochastic Neighbor Embedding (tSNE) visualization of 8 epithelial and 4 immune cell populations within the epithelium (n = 41,147 cells) from IMC data (n= 14 samples, 7 progressive, 7 regressive). (**B**) Heatmap of the average log-Z normalized expression within the epithelium of canonical markers (left) and the average log-Z stromal normalized expression within the epithelium of NLRC5 and its downstream targets (right) in identified cell types from (**A**). Bar plot shows the relative proportion of each cell type. (**C**) Bubble plot shows the differential composition of cell types in hyper-metaplasia and dysplasia as compared to normal epithelium. Color represents the compositional estimate, and dot size represents the log FDR value computed by sccomp. (**D**) t-SNE visualization of epithelial cell populations colored by NLRC5-high (dark blue) versus NLRC5-low (light blue) expression. (**E**) Violin plot of the log-Z stromal normalized expression of NLRC5 target genes (TAP1, PSMB9, and B2M) stratified by the NLRC5-high and NLRC5-low labels. (**F**) tSNE visualization of 8 immune and 3 stromal cell population in the stroma region (n = 46,254 cells) from IMC data (n= 14 samples, 7 progressive, 7 regressive). (**G**) Heatmap of the average log-Z normalized expression within the stroma of canonical markers (left) and the average log-Z stromal normalized expression within the epithelium of NLRC5 and its downstream targets (right) in identified cell types from (**F**). Bar plot shows the relative proportion of each cell type. (**H**) Violin plot of log-Z stromal normalized expression of NLRC5 in the epithelium (red) versus the stroma or non-epithelial tissue (yellow). (**I** and **J**) Cellular neighbors for each cell were defined based on a centroid distance within 30 μm. Cell-cell proximity score within each ROI (n = 24) is computed by the permutation test in imcRtools. Cyan bar represents p value < 0.05 whereas orange bars are not significant. (**I**) Bar plot showing the change in spatial proximity of immune cell populations to NLRC5-high versus NLRC5-low epithelial cells, averaged across all ROIs. (**J**) Bar plot showing the change in spatial proximity of CD8 T cells to NLRC5-high versus NLRC5-low epithelial populations, averaged across all ROIs. P values were determined by sccomp differential composition test for (**C**), the two-sided unpaired Wilcoxon test for (**E**) and (**H**), the two-sided paired Wilcoxon test for (**I**) and (**J**). * P <= 0.05, **P <= 0.01, ***P <= 0.001, ****P <= 0.0001.

The epithelial cells were dichotomized into cells with high or low NLRC5 expression based on the average of the log-z-score stromal normalized NLRC5 expression across ROIs (**Figure 5D, Extended Data Figure 7D, 7E).** The expression of NLRC5 downstream targets (B2M, PSMB9, and TAP1) were significantly lower in NLRC5-low versus NLRC5-high cells (p<0.001, two-sided Wilcoxon test, **Figure 5E**) and NLRC5 and its downstream targets were significantly correlated across epithelial cells (**r>0.6, p<0.001, Extended Data Figure 7F)**. The proportion of NLRC5-low epithelial cells was enriched among KRT5-low basal cells, MUC5B low secretory cells and MUC5B+ secretory cells, while the proportion of NLRC5-high epithelial cells was enriched among basal cells, differentiating basal cells, CEACAM5+ peri-goblets cells CEACAM5+SCGB1A1+ secretory cells, and FOXJ1+ ciliated cells (FDR<0.05, sccomp differential composition test, **Extended Data Figure 7G)**. These results suggest that protein expression of NLRC5 and 3 of its downstream targets are correlated in the epithelium and that NLRC5-high epithelial cells are enriched in intermediate state cell populations (e.g., CEACAM5+ peri-goblets cells CEACAM5+SCGB1A1+ secretory cells) that are increased in abundance in hyper-metaplastic and dysplastic regions.

### Basal cells with high NLRC5 protein expression were in closer spatial proximity to CD8 T cells

Next, we wanted to assess the spatial proximity of immune cell populations to epithelial cells with NLRC5-high versus low protein expression. Toward this goal, we clustered the 46,254 high quality cells in the non-epithelial tissue regions to identify 12 cell clusters (**Figure 5F**) based on expression of canonical cell markers (**Figure 5G**). Each cell cluster had cells from all samples, and we did not observe significant associations between cell cluster abundance and smoking status (**Extended Data Figure 8A, 8B**). NLRC5 expression was significantly lower in the non-epithelial tissue regions compared to the epithelial regions (p<0.001, Wilcoxon test, **Figure 5H**), suggesting that NLRC5 is more active within the epithelium. Given that some immune cells in non-epithelium regions are located near the epithelial basement membranes, we perform spatial proximity analysis using both epithelial and non-epithelial region immune cells. We tested the spatial cell-cell proximity of NLRC5-high and NLRC5-low epithelial cells to immune cells using the neighborhood permutation test (cell neighbors defined by a centroid distance=30 μm, n=500 permutations). We observed that NLRC5-high epithelial cells had closer spatial proximity to CD8 T cells whereas NLRC5-low epithelial cells had closer spatial proximity to B cells and macrophages (two-sided paired Wilcoxon test, p = 0.05, **Figure 5I, Extended Data Figure 9B, 9C**). Among the epithelial cell types, NLRC5-high basal cells had significantly closer spatial proximity to CD8 T cells than NLRC5-low basal cells (two-sided paired Wilcoxon test, p < 0.05, **Figure 5J, Extended Data Figure 9B, 9C**). The observation that NLRC5-high basal cells are near CD8 T cells suggests that these cells may be important mediators of a CD8 T cell response to dysplastic lesion development.

### Basal cells in progressive versus regressive PMLs have decreased NLRC5 expression in hyper-metaplasia and dysplasia regions

Given that NLRC5-high basal cells are near CD8 T cells, we wanted to test whether there were cell type compositional differences between progressive/persistent and regressive lesions, as previously we have shown the reduction of CD8 T cells in progressive lesions^10^. Differential cell composition analysis showed the proportion of CEACAM5+ peri-goblet cells are significantly higher in the epithelial tissue regions of progressive/persistent lesions compared with regressive lesions (FDR< 0.01, sccomp differential composition test, **Figure 6A, Extended Data Figure 7H)**. In the non-epithelial tissue regions, CD8 T cells, B cells and M2 (CD163+ only) macrophages are decreased, while VIM+ stromal cells and αSMA+VIM+ stromal cells are increased in progressive/persistent lesions compared to regressive lesions (FDR<0.05, sccomp differential composition test, **Figure 6B, Extended Data Figure 8C**). Basal cells were not decreased in abundance in progressive/persistent lesions, so we next investigated differences in NLRC5 expression. We found that NLRC5 protein levels are increased in epithelial regions of hyperplasia, metaplasia, and dysplasia compared to normal epithelium (p<0.001, linear mixed-effect model, **Figure 6C**) in regressive compared to progression/persistent lesions. As NLRC5 increases with increasing histologic severity, but increases less in progressive/persistent lesions, we wanted to test the interaction effect between histology and progression status within the epithelial cell types. We found that NLRC5 expression in the basal cells, basal/differentiating cells, K5 low basal cells, and MUC5B+ secretory cells is decreased in progressive/persistent lesions compared to regressive lesions in the hyper-metaplasia and dysplasia regions (p<0.05, linear mixed-effect model, **Figure 6D, Extended Data Figure 9B, 9C**). These analyses suggest that NLRC5 expression is decreased within basal cells in progressive/persistent compared with regressive PMLs and thus may be associated with the observed depletion of CD8 T cells.

**Figure 6.**
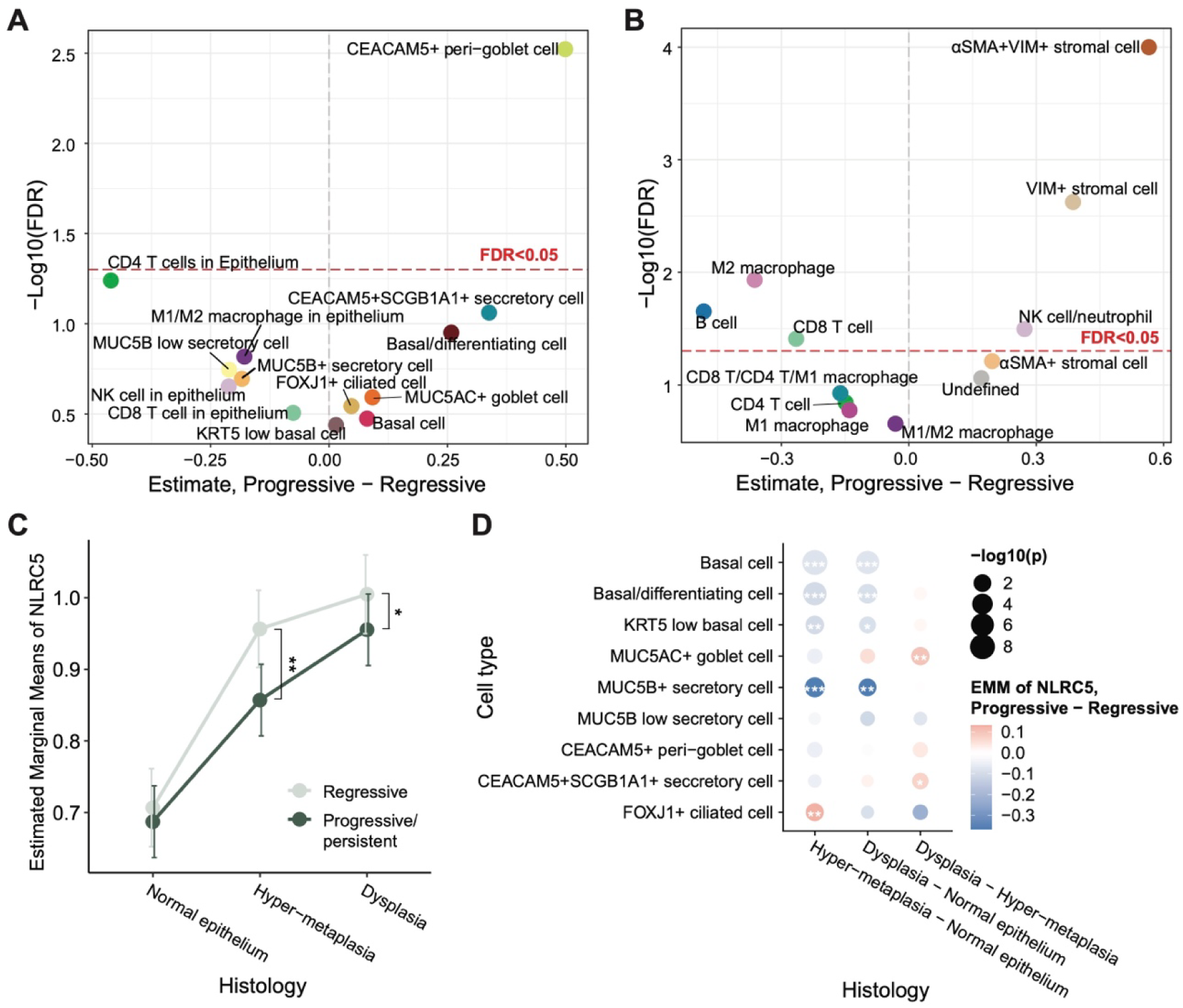
Immune and stroma cell composition and epithelial NLRC5 expression is associated with PML progression. Volcano plots showing the differential composition of progressive/persistent (n = 7) compared to regressive (n = 7) PMLs for (**A**) cell populations identified in epithelium and (**B**) cell populations identified in stroma regions. (**C**) Line plot showing the estimated marginal mean of NLRC5 expression with standard deviation in epithelial cells of progressive/persistent (dark green) and regressive (light green) PMLs, respectively, across the 3 histology regions. Although histology groups were modeled as categorical, the line plot is used to facilitate visualization of trends across histology grades. (**D**) Bubble plot showing the differential expression of NLRC5 between progressive/persistent and regressive PMLs in each epithelial cell population across the 3 histology contrasts. Color represents the estimate, and dot size represents the log p-value computed by linear mixed effect model. P values were determined by sccomp differential composition test for (**A**) and (**B**) and linear mixed effect models for (**C**) and (**D**). * P <= 0.05, **P <= 0.01, ***P <= 0.001.

## DISCUSSION

To move from treating to preventing lung cancer, a detailed understanding of the mechanisms that contribute to the development and progression of bronchial PMLs is needed. Our group and others have profiled the transcriptomes of bronchial PMLs, precursor lesions of LUSC, and showed that immune evasion is associated with increased histologic severity and lesion progression^6,10–12,36^. We previously demonstrated bronchial PMLs can be classified into four distinct molecular subtypes based on nine conserved gene co-expression modules. The proliferative molecular subtype was significantly enriched with bronchial dysplasia, and within this subtype, decreased expression of genes involved in antigen processing and presentation (module 9) and decreased immune cell infiltration are associated with persistent/progressive versus regressive PMLs^10^. In related prior work, progressive bronchial dysplasia was associated with decreases in T cells and HLA-DRA positive epithelial cells^12^. In progressive CIS lesions, genetic and epigenetic alterations in antigen presentation-related genes including loss of heterozygosity in the HLA region have been observed^11^. These findings support the hypothesis that impaired antigen processing and presentation contribute to lesion progression. However, the transcriptional regulators contributing to the dysregulation, especially in non-CIS PMLs, are still poorly understood. In this study, we measured miRNA expression profiles from longitudinally collected endobronchial biopsies previously profiled by mRNA sequencing and constructed a miRNA-mRNA module network to identify miRNAs associated with the antigen presentation gene co-expression module (module 9). We found that hsa-miR-149-5p, a miRNA highly expressed in the epithelium, was up-regulated among the persistent/progressive compared to regressive proliferative PMLs and repressed NLRC5, the transcriptional regulator of MHC-I genes. Furthermore, we used miR-ISH and imaging mass cytometry to characterize the spatial localization of hsa-miR-149-5p, NLRC5, and its downstream targets in the context of the lesion cellular architecture and the immune microenvironment in biopsy tissue sections. Our results demonstrate that NLRC5 and the MHC-I genes that it regulates are increased with increasing histologic severity in the epithelium. However, when comparing progressive/persistent versus regressive proliferative subtype PMLs within different histologic grades, there is a decrease in expression of NLRC5 and its downstream targets partially driven by increased hsa-miR-149-5p expression. Decreased NLRC5 in basal cells was associated with reduced spatial proximity to CD8 T cells and may contribute to the immune evasion phenotype.

Among the direct targets of hsa-miR-149-5p within Module 9 was NLRC5, a gene that regulates the expression of MHC-I (HLA-A, HLA-B, HLA-C, B2M) and immunoproteasome genes (PSMB8, PSMB9, TAP1). miR-ISH analysis indicated that hsa-miR-149-5p was expressed in epithelial cells of PMLs and *in vitro* experiments showed NLRC5 and several of its targets were decreased by hsa-miR-149-5p overexpression in a human lung epithelial cell line. Several studies, in different cancer contexts, support the oncogenic role of increased hsa-miR-149-5p^21^. hsa-miR-149-5p is up-regulated in non-small cell lung tumors compared with adjacent tissues^37^, lower expression of hsa-miR-149-5p inhibited cell growth in prostate cancer cells *in vitro*^38^, and higher exosomal hsa-miR-149-5p was associated with metastatic melanoma^39^. Notably, Srivastava *et al*. showed that IFN-γ activity may suppress hsa-miR-149-5p levels and drive an inflammatory response in keratinocytes^40^. We observed a similar association where progressive/persistent proliferative PMLs have decreased expression of interferon signaling genes and high levels of hsa-miR-149-5p compared with regressive PMLs. Additionally, defects in antigen processing and presentation are hallmarks of cancer immune evasion^41^. Many of the targets of hsa-miR-149-5p are inactivated by other oncogenic mechanisms to alter the antigen presentation activity in cancer cells, including the loss of heterozygosity of HLA and B2M genes^42,43^, somatic mutation in HLA genes^44^, epigenetic alterations^45,46^, and transcriptional regulation^30^. Less is known in bronchial PMLs, however, recent work showed that the loss of heterozygosity and hypermethylation in the HLA regions are associated with dysfunctional antigen processing and presentation in CIS lesions^11^. Our findings suggest a novel post-transcriptional regulatory mechanism by which up-regulation of hsa-miR-149-5p in airway basal cells impairs antigen processing and presentation contributing to immune evasion.

Demonstrated by miR-ISH, hsa-miR-149-5p expression is increased in epithelial tissue regions compared to non-epithelial regions, and in regions of dysplasia, its expression is higher in progressive/persistent compared to regressive PMLs. The resolution of hsa-miR-149-5p expression to basal cells, however, was not definitive as we did not have a KRT5 counter stain. In the IMC data, we observed a significant decrease in NLRC5 expression in basal, basal/differentiating, K5 low basal, and MUC5B+ secretory cells in regions of hyper-metaplasia and dysplasia in progressive/persistent versus regressive PMLs. Since hsa-miR-149-5p expression is assayed across the whole tissue via miR-ISH, but the IMC results are based on regions of interest, direct integration of the results was not possible. The results, however, suggest that hsa-miR-149-5p and NLRC5 expression were anti-correlated in regions of dysplasia. Out of the epithelial cell types with decreased NLRC5 expression associated with progression, basal cells were the only cell type where the level of NLRC5 influenced its spatial proximity to CD8+ T cells. Basal cells are the progenitor stem cell and suggested to be the cell of origin of LUSC in the proximal airway^47,48^. Laughney *et al*. demonstrated an inverse association between expression levels of SOX2 (a transcription factor specifying lung cell fate) and MHC-I genes in primary and metastatic lung tumors^49^. Similarly, the stem cell program in colon cancer has been associated with decreased levels of antigen presentation^50^. These observations suggest that the cell states of basal cells may be an important determinant of the immune surveillance and evasion in bronchial PML progression. Interestingly, we also observed that CEACAM5+ peri-goblet cells, a smoking associated cell population^33^, were enriched with increasing histologic severity and in progressive/persistent compared to regressive Proliferative PMLs. We did not, however, detect a change in NLRC5 expression in these cells associated with progression/regression status or that NLRC5 levels influence their spatial proximity to CD8+ T cells. Future investigations will be needed to further unravel the causality between epithelial cell state, expression of hsa-miR-149-5p and NLRC5, and the functional ability of cells to present antigen and recruit immune cells.

The IMC data also revealed that NLRC5 protein expression increases as the histology grade advances from normal epithelium to hyper-metaplasia and dysplasia accompanied by an increase in CD8 T cells, CD4 T cells and NK cells. In progressive/persistent compared to regressive proliferative PMLs there is a decrease in NLRC5 protein expression within hyper-metaplasia and dysplasia regions and CD8 T cells, B cells, and M2 macrophages are decreased. Spatial analysis demonstrated that NLRC5-high basal cells are in significantly closer proximity to CD8+ T cells compared to NLRC5-low basal cells. Levels of NLRC5 were also shown to decrease in basal cell populations in areas of hyper-metaplasia and dysplasia in progressive/persistent compared to regressive Proliferative PMLs. Similar relationships between NLRC5 regulated MHC-I gene expression and cancer immune evasion have been extensively studied in the context of cancer^29,30^. Using the solid tumor data from TCGA, Yoshihama *et al*. demonstrated that lower NLRC5 expression is associated with deficient CD8+ T cell activation and poor prognosis^51^. Furthermore, recent work demonstrated that impaired MHC-I processing and presentation pathway is related to resistance to immune therapy in lung cancer^52^, and increased MHC-I gene expression can improve immunotherapy response^53^, suggesting their critical roles and therapeutic potential in the context of lung cancer management.

In summary, utilizing miRNA-gene network with miR-ISH and imaging mass cytometry assays to identify and resolve the spatial and cellular localization of hsa-miR-149-5p, its downstream targets, and the immune microenvironment we have identified a novel post-transcriptional regulatory mechanism that may contributing to immune evasion and bronchial PML progression. Our results suggest cell type specific biomarkers of lesion progression and a potential epithelial-specific candidate therapeutic target to potentially reverse early immune evasion. Future experiments that perturb hsa-miR-149-5p, NLRC5, and MHC-I gene expression in model systems are required to confirm the extent to which hsa-miR-149-5p regulates NLRC5 levels and the immune microenvironment and the association between MHC-I gene expression and recruitment of cytotoxic CD8 T cells. Additionally, testing anti-mir oligonucleotides^54,55^ to target hsa-miR-149-5p in preclinical models such as the NTCU mouse mouse^56,57^ are needed to show its efficacy in preventing or delaying lesion progression. Uncovering potential regulators of early immune evasion holds promise to advance lung cancer chemoprevention in patients at high-risk for developing lung cancer.

## METHODS

### Sample collection

The patient enrollment and sample collection procedures were described previously^10^. The study was conducted with approval from the Roswell Park Comprehensive Cancer Center and the Boston University School of Medicine Institutional Review Boards and all subjects provided written informed consent. Briefly, endobronchial biopsies and bronchial brushings were collected via autofluorescence bronchoscopy from high-risk individuals in a low-dose computed tomography lung cancer screening program at approximately 1-year intervals. Biopsy outcome status was defined as previously described^10^ where the outcome status of each biopsy (progressive/persistent, regressive, normal stable, or unknown) was based on the worst histology recorded at the same anatomic location in future procedures. During the bronchoscopy, two biopsy samples were obtained from each sampled lung anatomic location. One biopsy was formalin fixed and paraffin embedded (FFPE) and used for histological evaluation, miRNA in situ hybridization (miR-ISH), and image mass cytometry (IMC) assays, and the other biopsy was frozen and used for RNA isolation and mRNA and miRNA sequencing. Bronchial brushes were collected at each bronchoscopy from a normal appearing area of the right or left mainstem bronchus. In our prior study^10^, patients were divided into a Discovery (n=197 biopsies, n=91 brushes, n=30 patients) and Validation Set (n=111 biopsies, n=49 brushes, n=20 patients) based on time of initial sample collection. In this study, Discovery cohort samples with sufficient miRNA underwent miRNA library preparation and sequencing (167 biopsies and 91 brushes, n=30 patients). Additionally adjacent fixed samples (19 biopsies, n=9 patients) were subject to detailed H&E annotation, miR-ISH and IMC assays (**Figure 1A**).

### miRNA library preparation, sequencing, and data pre-processing

miRNA was extracted from biopsies or brushing samples using the miRNeasy Mini kit (Qiagen) according to the manufacturer’s instructions^10^. miRNA sequencing libraries were prepared using NEBNext Multiplex Small RNA Library Prep Set for Illumina (NEBioLabs). Sequencing adapters that target the 3’ hydroxyl group of small RNAs were ligated and the transcripts were reverse transcribed and PCR-amplified into single-stranded cDNA libraries. The libraries were pooled, and size selected for small RNA fragments on PAGE gel in groups of 6-10. The libraries were sequenced on Illumina® HiSeq 2500 to generate more than 10 million single-read 35-36 nucleotide (nt) reads per sample^58^.

De-multiplexing and generation of FASTQ files were performed using Illumina CASAVA v1.8.2. FastQC v0.11.7 was used to examine the quality of raw reads and cutadapt v1.18 was used for trimming the sequencing adaptors (5’- AGATCGGAAGAGCACACGTCTGAACTCCAGTCAC-3’, -trim-n)^59,60^. Reads unlikely to be properly sequenced mature miRNA transcripts were discarded (reads shorter than 16 nt or longer than 25 nt with –m 16 –M 25 in cutadapt). Alignment and quantification of mature miRNAs based on miRBase v22 were performed with miRDeep2 v0.1.0^61,62^ with mapper.pl parameters –e –h –m –i –j and quantifier.pl parameter –k –j –d -P. For mature miRNAs associated with multiple precursor miRNAs, the max counts were used.

We performed miRNA-seq sample quality assessments and miRNA filtering separately for each sample type (endobronchial biopsies and bronchial brushes). We limited the analysis to the miRNA samples with previously analyzed (GSE109743, dbGaP:phs003185.v1.p1) matched mRNA expression profiles (n=160 biopsies, n=89 brushes)^10^. The raw counts were converted into log2 counts per million of miRNA reads mapped (logCPM) and quantile normalized with voom^63^. miRNAs with logCPM less than 1 in more than half of the samples were removed. We excluded samples (n=4 biopsies, n=2 brushes) if sequencing quality metrics were outside 2 standard deviations from the mean. The quality metrics we used included the first and the second principal components from the PCA using filtered miRNA expression, the mean Pearson expression correlation with all other samples across the filtered miRNAs, and the transcript integrity number (TIN) of the matched mRNA sequencing sample (calculated using RSeQC v3.0.0^64^). Finally, 156 biopsies and 87 brush samples from 30 patients were retained. The gene filter and normalization described above were performed on the final high-quality samples with matching mRNA data (n=525 miRNAs in biopsies and n=489 miRNAs in brushes). The residual miRNA expressions adjusting for sequencing batch were computed with limma^65^ for downstream analysis.

### miRNA-gene network construction to identify miRNAs associated with gene co-expression modules that define PML molecular subtypes

To capture the association of miRNAs with the previously described^10^ 9 co-expression gene modules within biopsy samples, we first constructed a miRNA-gene regulatory network focused on the module genes (n=3936; **Supplementary Table 2**)^66^. For this network, we utilized both the predicted gene targets for a miRNA and the correlation between a miRNA and its targets (**Figure 1B**). The predicted gene targets for each miRNA were defined as those identified by any of the three miRNA target databases, including TargetScan v7.2, starBase, and miRTarBase v7.0^26–28^. Between each miRNA and its predicted target gene, the Pearson correlation coefficient was calculated using the residual expression values, of miRNA (described above) and gene (adjusted for median TIN and batch as in Beane *et al* ^10^), and the significance level was adjusted using false discovery rate (FDR). Only the edges with significant and negative Pearson correlation coefficients (FDR <= 0.05) between miRNA and predicted target genes were retained in the network.

After constructing the miRNA-gene regulatory network, we sought to retain only connections where miRNAs had regulatory relationships with a gene module that were stronger and more specific than with other gene modules (**Figure 1B**). For each miRNA, we first performed Fisher-Z transformation on the Pearson correlation coefficient density of a miRNA with its targets and compared to the density with its targets within the module to the density with targets in other modules using a two-sample or one-sample (if only one target is within a group) t-test to select miRNAs with strong negative relationships to modules. We also used Fisher-exact tests to determine whether a miRNA had more significantly negative-correlated target genes in the module of interest than in other modules (odds ratio > 1). The results of both tests were adjusted using FDR and miRNAs with significant results for the same gene module from both tests (FDR <= 0.05) were identified as specifically regulating that module. Any connection between a miRNA and predicted target genes of a module that did not pass both tests was removed. The final network can be viewed as a filtered subset of the full miRNA-gene regulatory network that captures the potential regulatory relationship between miRNAs and gene modules (**Figure 2A**). Each miRNA in the network was assigned to its associated one or more modules (module-associated miRNA) Summary scores were computed based on miRNAs connected to each gene module^67^ using Gene-set Variation Analysis (GSVA).

### Identification of miRNAs and genes associated with PML progression status

The association between miRNAs that may regulate gene modules and PML progression status was determined based on whether the expression level of a miRNA is significantly different between the progressive/persistent and the regressive PMLs of the proliferative subtype (p-value < 0.05). Within each model, miRNA residual expressions were used as the dependent variable, PML progression status was the main independent variable, and patient was included as a random effect using ‘duplicateCorrelation()’ function from limma v3.44.3^65^.

To identify the genes that are associated with PML progression status, we used the full gene expression data of the discovery and validation cohorts from our previous publication, including those samples without matched miRNA sequencing data^10^. Similar linear mixed-effect models were performed with gene residual expressions (adjusted for sequencing batch and TIN) as the dependent variable, PML progression status as the independent variable while adjusting patient as a random effect. We further validated gene associations with PML progression status using an external microarray dataset^12^ (GSE114489), in which linear models were used with probe intensities as the dependent variable and progression status as the main independent variable. The progressive/persistent group included the persistent bronchial dysplasia and progressive non-dysplasia samples based on the original annotation, and the regressive group included the regressive bronchial dysplasia group.

The association between hsa-miR-149-5p and the antigen presentation gene module was examined in three datasets with sample-matched miRNA and gene expression profiles, including the airway brushing samples from patience with PMLs, TCGA LUSC samples, and AEGIS samples. Gene set enrichment analysis (GSEA)^68^ was performed with R package fgsea^69^ with random seed of 1234 and nperm=10000, where genes were ranked by their Pearson correlation coefficient with hsa-miR-149-5p. The negative correlation between hsa-miR-149-5p and NLRC5 was also validated in these three datasets using Pearson correlation.

### Examination of the cell type-specific expression of hsa-miR-149-5p

To examine whether a miRNA is potentially specifically expressed within a certain cell type, we examined the correlation between the expression level of the miRNA and cell-type marker genes. The method for calculating Pearson correlation coefficients has been described in the previous section. The following cell type marker genes were used: *CD3G* for T cells, *CD19* for B cells, *CD68* for macrophages, *KRT5* for basal cells, *FOXJ1* for ciliated cells, *MUC5AC* for goblet epithelial cells, *SCGB1A1* for club cells, *MUC5B* for mucus secretory cells and *CEACAM5* for peri-goblet cells^33^. Also, miRNA expression data from functional annotation of the mammalian genome 5 (FANTOM5) project were used to evaluate whether the expression of a miRNA is enriched within a certain group of samples^43^. Classification of FANTOM5 samples to cell types were based on cell ontology annotation from FANTOM5. Samples belonging to the “epithelial cell” or “leukocyte” group were used as the epithelial and immune compartment. Samples were ranked by their normalized expression levels of hsa-miR-149-5p and GSEA^68^ using R package fgsea^69^ with seed 1234 and nperm=10000 was used to test whether samples with higher hsa-miR-149-5p expression were enriched for samples from a specific group of cell types. Gene expression data from FANTOM5 were also used to examine the correlation between hsa-miR-149-5p and *NLRC5* among different cell-types.

### In vitro perturbation hsa-miR-149-5p and *NLRC5* expression

Human LUSC cells, SW900 (ATCC), were grown in RPMI medium and transfected using Lipofectamine RNAiMAX (Thermo) following the manufacturer’s protocol. Cells were transfected with 1) hsa-miR-149-5p (*mir*Vana, Thermo); 2) miR scramble control (MISSION, Sigma); 3) siNLRC5 (NLRC5 84166 siRNA, SMARTpool, Dharmacon). Cells were incubated in transfection mix for 48 hours, washed, and lysed in Qiazol (Qiagen) for 5 min at room temperature before being stored at -80°C until RNA isolation. Following RNA isolation, qRT-PCR was conducted using 1 ng of RNA per sample that was reverse transcribed using the Transcriptor First Strand cDNA Synthesis Kit (Roche) following the supplieŕs protocol. For miRNA quantification, 1 ng total RNA was used in cDNA synthesis via the TaqMan Advanced miRNA cDNA Synthesis Kit (Thermo). qRT-PCR was performed using the TaqManTM Fast Advanced Master Mix (Applied Biosystems) and TaqMan probes (Thermo): *PPIA* (Hs04194521_s1), *NLRC5* (Hs01072123_m1), *HLA-A* (Hs01058806_g1), *HLA-B* (Hs00818803_g1), *HLA-C* (Hs00740298_g1), *PSMB9 (*Hs00160610_m1), *TAP1* (Hs00388675_m1), hsa-miR-149-5p (477917_mir), and hsa-miR-98-5p (478590_mir). The 2-ΔΔCT method^70^ was used for relative transcript quantification. *PPIA* was used as the reference gene for mRNA quantification. For miRNA quantification, miR98 served as an internal control. The average ΔCT of mock controls in each 96-well qPCR plate (representing experiment batch) was used as the reference sample. Gene expression differences between treatment conditions were determined by two-sided Wilcoxon test.

### Biopsy selection, sectioning, staining, and pathologist annotation of digitized images

FFPE tissues sections from 19 proliferative biopsies from 9 patients (n=10 progressive/persistent and n=9 regressive) were selected from samples where adjacent biopsies were profiled by mRNA-seq and miRNA-seq (**Supplemental Table 8**). Three sequential tissue sections were stained using H&E for annotation of histopathology features, chromogenic miRNA in situ hybridization (miRNA-ISH using miRNAscope) with a hsa-miR-149-5p probe to assess hsa-miR-149-5p expression, and imaging mass cytometry (IMC) using a cocktail of 28 antibodies targeted to epithelial and immune populations as well as NLRC5 and its downstream targets (for 14 out of the 19 biopsies). The H&Es were scanned at 40X using the Leica Aperio AT2 scanner and the miRNA-ISH slides were scanned under oil immersion at 63X using the Leica Aperio Versa Scanner. The digitized H&E image files were uploaded to the deepPath Pixelview software (https://www.deeppath.ai) and the epithelial tissue was annotated by the pathologist (E.J. B.) by creating masks for the following: normal epithelium, reserve cell hyperplasia, goblet cell hyperplasia, immature squamous metaplasia, squamous metaplasia, and mild, moderate, or severe squamous dysplasia. These annotations were manually transferred to digitized images of the miRNA-ISH and IMC using QuPath v0.3.2 software^71^ and reviewed for accuracy by the pathologist.

### Quantification of hsa-miR-149-5p expression in biopsies via miRNAscope

For miRNA-ISH, 5 µm FFPE tissue sections were stained using the miRNAscope HD Assay Red (324530, ACD Bio, Newark, CA) with probe SR-hsa-miR-149-5p-S1 (1058711-S1, ACD Bio, Newark, CA). The staining was performed using the manufacturer’s protocol, however, treatment with Protease III was reduced to 10 min to conserve tissue morphology of the small biopsy samples.

A pixel classifier to detect the spots of hsa-miR-149-5p signal was trained on a representative region (250000 µm^2^) selected in each sample using the artificial neural network (ANN) model from QuPath v0.3.2 software with default multiscale features of 3 channels – hematoxylin, probe stain, and residual channels^71^. To validate the classifier performance, we selected a representative region (16726 +/- 4332 mm^2^) different from the training regions in each sample and annotated the hsa-miR-149-5p spots manually. The recall (calculated as true positive (TP) divided by TP plus false negative (FN)), the precision (calculated as TP divided by TP plus false positive), and the F1 score (calculated as the harmonic mean of recall and precision) were calculated per sample. The hsa-miR-149-5p density is defined either as hsa-miR-149-5p predicted spots per annotated area or as hsa-miR-149-5p predicted spots per cell. The cell number was estimated by nuclei segmentation using StarDist extension v0.3.2^72^ with the pre-trained brightfield model implemented via QuPath software in regions annotated as epithelium. The association of hsa-miR-149-5p density with progression status was modeled as a linear mixed-effect model, with the interaction of progression status and histology as fixed effect and sample ID as random effect using lmerTest v3.1-3^73^.

### Imaging mass cytometry (IMC) staining and ablation

The IMC experiments assayed the levels of 28 proteins and DNA (**Supplementary Table 10**). Antibodies were pre-conjugated (Standard BioTools) or conjugated to lanthanide isotopes following the MaxPar Antibody Labeling kit manufacturers’ protocol. The IMC staining was done using the proximal level of the 14 FFPE tissue sample slides analyzed via miRNAscope^TM^ assay (**Supplementary Table 8**). The slides were dewaxed in xylene for 10 min, followed by two more xylene washes of 5 min each. Tissue was rehydrated in sequential alcohol dilutions of 100% (three consecutive washes), 95% (two consecutive washes), 80% and 70% for 10 min each before washing in deionized water for 10 min. Heat induced epitope retrieval was performed in Tris-EDTA pH8.5 antigen retrieval buffer containing 5% glycerol and heating to 96°C in the microwave for 30 min. Slides were allowed to cool and washed twice in deionized water and two washes in PBS (10 min each). Samples were blocked with 3% BSA in PBS for 45 min at room temperature in a wet chamber. The antibody cocktail was prepared to a final concentration of 0.5% BSA and added to the samples for overnight incubation at 4°C. The stained samples were washed twice in 0.2% TritonX100/PBS followed by one wash in PBS for 8 min each. A 1:1000 dilution of Iridium intercalators (Standard BioTools) in PBS were used to stain DNA for 30 min at room temperature. Finally, slides were washed in PBS for 8 min and deionized water for 5 min before air drying. Regions of interest (ROIs) were laser ablated using the Hyperion Imaging Mass Cytometry^TM^ system with a 200 Hz frequency at 1 µm^2^ resolution.

### IMC data processing and quality control

The IMC data was segmented into single cells and single cell data was extracted using the Steinbock (v0.14.1) toolkit^74^. Briefly, the raw data (in MCD format) were converted to standardized OME-TIFF files and single cell masks were generated using the TissueNet model in Cellpose^75^. DNA1 and DNA2 were used as the nuclei channel, and multiple markers, including KRT5, KRT8, E-Cadherin, SCGB1A1, MUC5AC, MUC5B, FOXJ1, CEACAM5, CD3, CD4, CD8A, CD20, CD68, HLA-DR, αSMA, and Vimentin (VIM), were used as the membrane/cytosol channel. Single cell features, including mean intensity of each marker and spatial coordinates, were extracted from the single-cell masks across all available channels. The extracted single cell data was converted to a SpatialExperiment object using the imcRtools v1.8.0 R package^74^ for downstream analysis. Regions of interest (ROIs) less than 1000 cells were removed from this analysis (n=7 ROIs) and cells with an area less than 1st percentile (21 µm^2^) or larger than 98th percentile (198 µm^2^) were filtered out (n=2804 cells). A total of 87,401 cells from 24 ROIs were used for final analysis. Antibodies with poor signal or non-specific staining (HLA-ABC, Ki-67, CD25) were excluded from the analysis.

### IMC data clustering and differential composition analysis

The mean intensity of the single-cell raw measurement for each marker was log-transformed, and z-score normalized with respect to each ROI, referred to as log-Z normalization. The canonical phenotypic markers for epithelial cells (KRT5, KRT8, p40, E-Cadherin, MUC5AC, MUC5B, CEACAM5, SCGB1A1, FOXJ1), immune cells (CD20, CD3, CD4, CD8A, GZMB, PDL1, CD68, CD163, HLA-DR), and stromal cells (αSMA, VIM) were used for UMAP dimension reduction, fastMNN batch correction^76^ and PhenoGraph clustering^77^ of all the cells. Broad cell types, including epithelial, immune, and stromal cells, were annotated based on the expression of KRT5, E-Cadherin, CD20, CD3, HLA-DR, αSMA, and VIM. Given that the proportions of epithelial and non-epithelial regions are different across ROIs, we analyzed the epithelial and non-epithelial (including broad immune and stromal cells) regions separately. For each region type (epithelial or non-epithelial), signals were log-Z normalized within each ROI and used for tSNE dimension reduction and PhenoGraph clustering^77^. In the non-epithelium compartment, a cluster was identified where more than half of the cells have epithelial histology annotations. 1260 cells with any epithelial histology annotation in this cluster, which are shown to localized in epithelium in IMC images, are reclassified as broad epithelial cells. Then, we repeated the clustering analysis as described above. Cell types in epithelial regions were identified based on epithelial and immune cell markers, while non-epithelial cell types were identified based on immune and stromal cell markers. The differential composition analysis of the cell types by histology, smoking, and progression status was performed using sccomp v.1.7.14^78^ within each region type.

### Quantification of NLRC5 protein expression and its association with tissue and sample characteristics

We observed that the NLRC5 levels were significantly higher for cells within epithelium compared to the non-epithelial regions. To compare the NLRC5 levels across ROIs, we resampled the cells in non-epithelial regions to equalize the proportions of cells between epithelial and non-epithelial regions in each ROI, followed by z-score normalization of the log-transformed NLRC5 intensities, referred to as log-Z stromal normalized NLRC5. Epithelial cells were dichotomized as NLRC5-high or NLRC5-low using the overall average of the log-Z stromal normalized NLRC5 across all epithelial cells as the cut off. The association of log-Z stromal normalized NLRC5 expression with progression status was modeled using a linear mixed-effect model, with the interaction of progression status and histology as fixed effect and ROI as random effect using lmerTest v3.1-3^73^. The estimated marginal means for the fitted model were computed by emmeans v1.10.3^79^.

### IMC spatial cell-cell proximity analysis

Cell neighbors for each cell were identified by distance-based expansion with a centroid distance threshold of 30 mm using buildSpatialGraph() function in imcRtools v1.8.0^74^. Immune cell type annotations from both epithelial and non-epithelial regions were merged for the same cell types. Broad epithelial or epithelial subtypes are stratified by the NLRC5 level as previously described. Spatial cell-cell proximity analysis for each ROI was performed using standard neighborhood permutation test^80^ with 500 iterations using testInteractions() function in imcRtools v1.8.0^74^. The spatial proximity score was defined as the fraction of perturbations having proximity counts equal to or less than the observed counts for each pair of cell type entries. Difference in spatial proximity between NLRC5-high and NLRC5-low epithelial cells or epithelial subtypes with respect to immune subtypes were assessed by the two-sided paired Wilcoxon test.

### Data availability

#### Endobronchial Biopsy and Airway Brushing miRNA Expression Data

Raw FASTQ files and the miRNA expression matrices and associated sample and clinical data are available on the Human Tumor Atlas Network Data Portal (https://humantumoratlas.org) using the identifications provided in **Supplemental Table 11.**

#### Endobronchial Biopsy Imaging Data

H&E, miR-ISH, and IMC images are also available on the Human Tumor Atlas Network Data Portal (https://humantumoratlas.org) using the identifications provided in **Supplemental Table 11.**

#### Endobronchial Biopsy and Airway Brushing Gene Expression Data

Raw FASTQ files are available via dbGaP accession phs003185.v1.p1 and the gene expression count and residual matrices are available via Gene Expression Omnibus accession number GSE109743. Gene expression residual data from GSE109743 was used to construct miRNA-mRNA correlation network^10^.

#### Additional Publicly Available Endobronchial Biopsy Data

miRNA expression data from GSE93284 and gene expression from GSE66499 and GSE114489 were used to validate the miRNA-mRNA correlation and gene association with lesion outcome^12,23,81^.

#### FANTOM5 Gene & miRNA Expression Data

*FANTOM5* miRNA expression profiles for 2595 mature miRNAs across 399 samples (directly downloaded from https://fantom.gsc.riken.jp/5/suppl/De_Rie_et_al_2017/ in November 2022)^35^. The library size normalized (CPM) miRNA expression matrix was log2-transformed with pseudocount of 0.1. Matched gene expression profiles in transcripts per million (TPM; N=394) were downloaded from the same website and were used to examine the miRNA-mRNA correlation by cell type. and the tag count of the most supported peak (p1@NLRC5) was used for the expression of NLRC5 per sample. The TPM values were log2-transformed with pseudocount of 0.1 for correlation analysis.

#### TCGA-LUSC mRNA & miRNA Expression Data

Sample matched gene and miRNA expression data for the TCGA LUSC (N=475) were also used to validate the miRNA-mRNA correlation. Gene (legacy) and miRNA expression data of the primary tumor samples were obtained using TCGAbiolink^22^ in October 2022. Counts associated with the same mature miRNA transcript were aggregated. miRNA counts normalization was performed as described above. The Pearson correlation coefficients were then calculated between miRNA and mRNA data on expression residuals adjusting for the plate as previously described^10^.

#### Gene and miRNA expression from bronchial brushings from the Airway Epithelial Gene Expression in the Diagnosis of Lung Cancer (AEGIS) trials

Gene expression and miRNA expression data from sample matched mainstem bronchus brushing samples (N=341) were obtained from AEGIS clinical trials. Gene-level expression data was obtained from GSE66499^24^ and was processed as previously described^82^. miRNA sequencing data was obtained from GSE93284^23^.

#### miRNA Target Database

Three target databases for miRNA target information were used. For TargetScan v7.2^26^, all conserved miRNA sites were downloaded from targetscan.org. miRNAs with an additional suffix (.1 or .2) compared to miRbase were both included. For starBase^27^, all predictions for hg19 were downloaded from starbase.sysu.edu.cn on December 2022 using the API and were filtered for clipExpNum >= 5. For miRTarBase v7.0^28^, the HGNC symbols were translated to Ensembl IDs (GRCh37) using biomaRt^83^.

### Code availability

All code will be made available upon publication via the following Github repository, https://github.com/jbeane0428/miR149.git.

