## Extended Data Figures for "Epithelial miR-149-5p up-regulation is associated with immune evasion in progressive bronchial premalignant lesions"

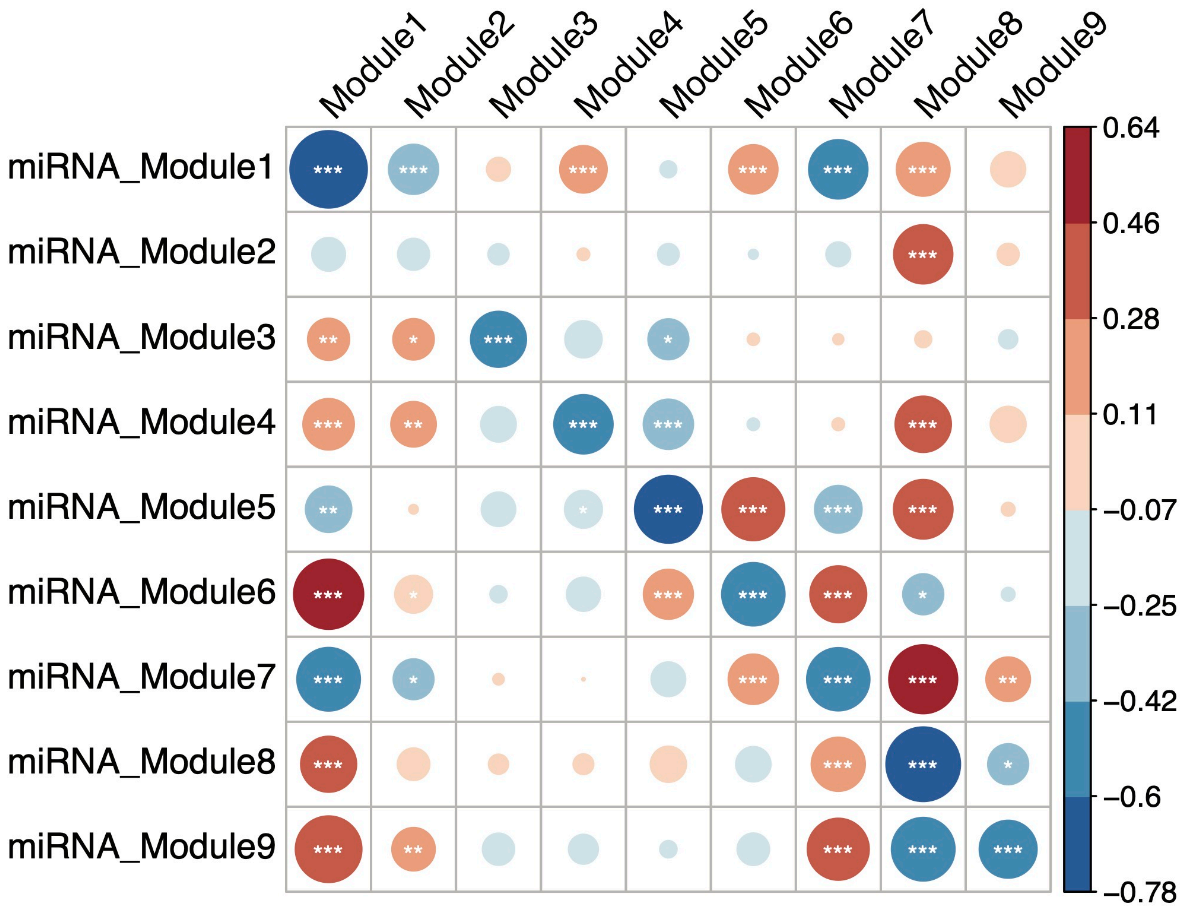


**Extended Data Figure 1. Correlation between GSVA scores of miRNAs and gene modules from the miRNA-gene network.** Bubble plots of the Pearson correlation between GSVA scores of module associated miRNAs and gene modules from the miRNA-gene module network. Each miRNA connected to predicted target genes within a gene module and that has passed the statistical tests outlined in the methods was assigned to that gene module. The expression value of all miRNAs assigned to a gene module was calculated using GSVA. * FDR <= 0.05; ** FDR <= 0.01; *** FDR <= 0.001.

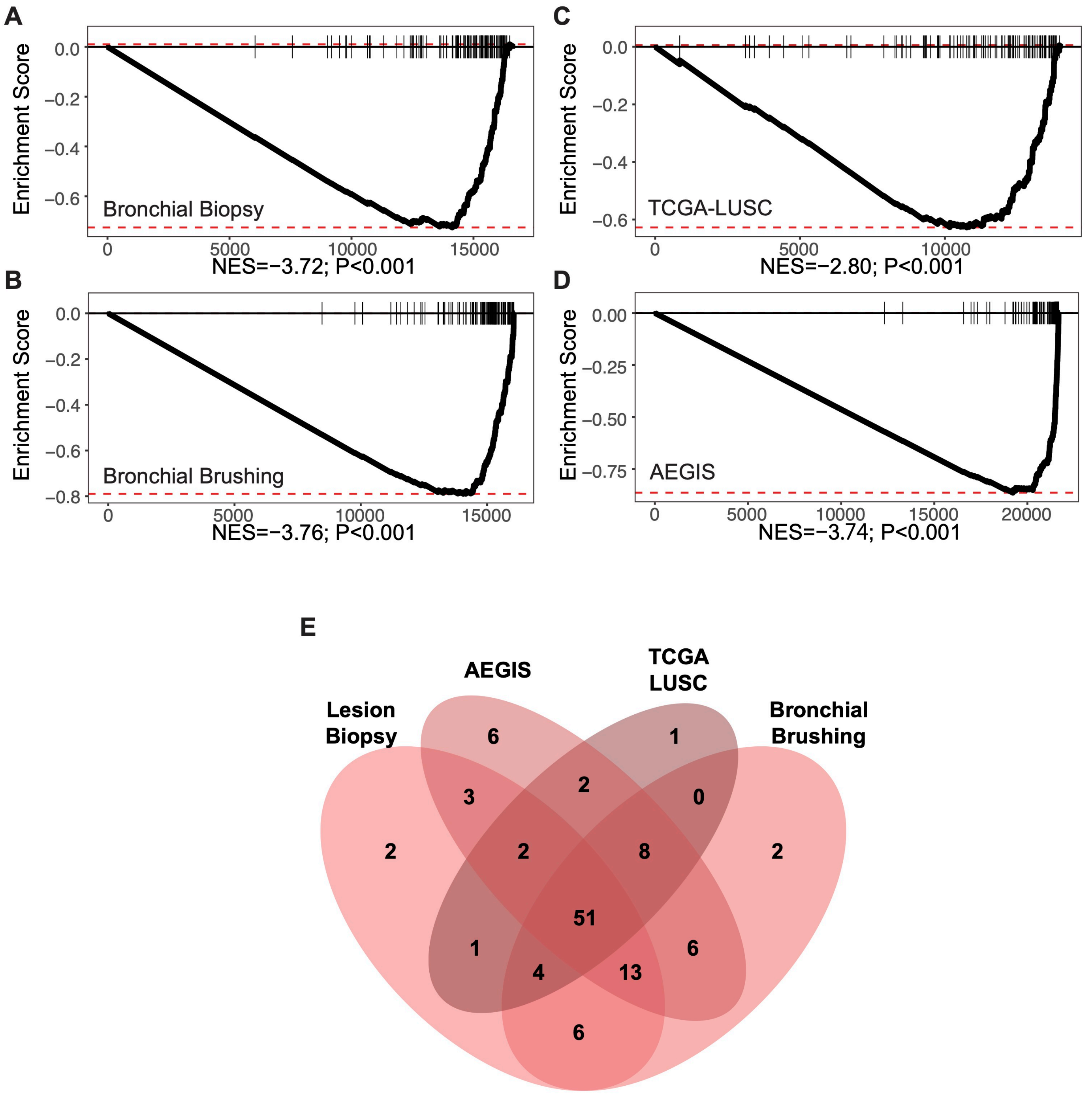


**Extended Data Figure 2. Antigen presentation module genes are enriched among genes negatively correlated with hsa-miR-149-5p in lung-related datasets.** Enrichment plot of module 9 genes (n=112) among all genes ranked by their expression level correlation with hsa-miR-149-5p across four datasets: (**A**) biopsy samples (n=156), (**B**) bronchial brushing samples (n=87) from this study, (**C)** TCGA-LUSC primary tumor samples (n=475), (**D**) AEGIS bronchial brushing samples (n=341). (**E**) Overlap of leading-edge genes from analyses in **A-D**.

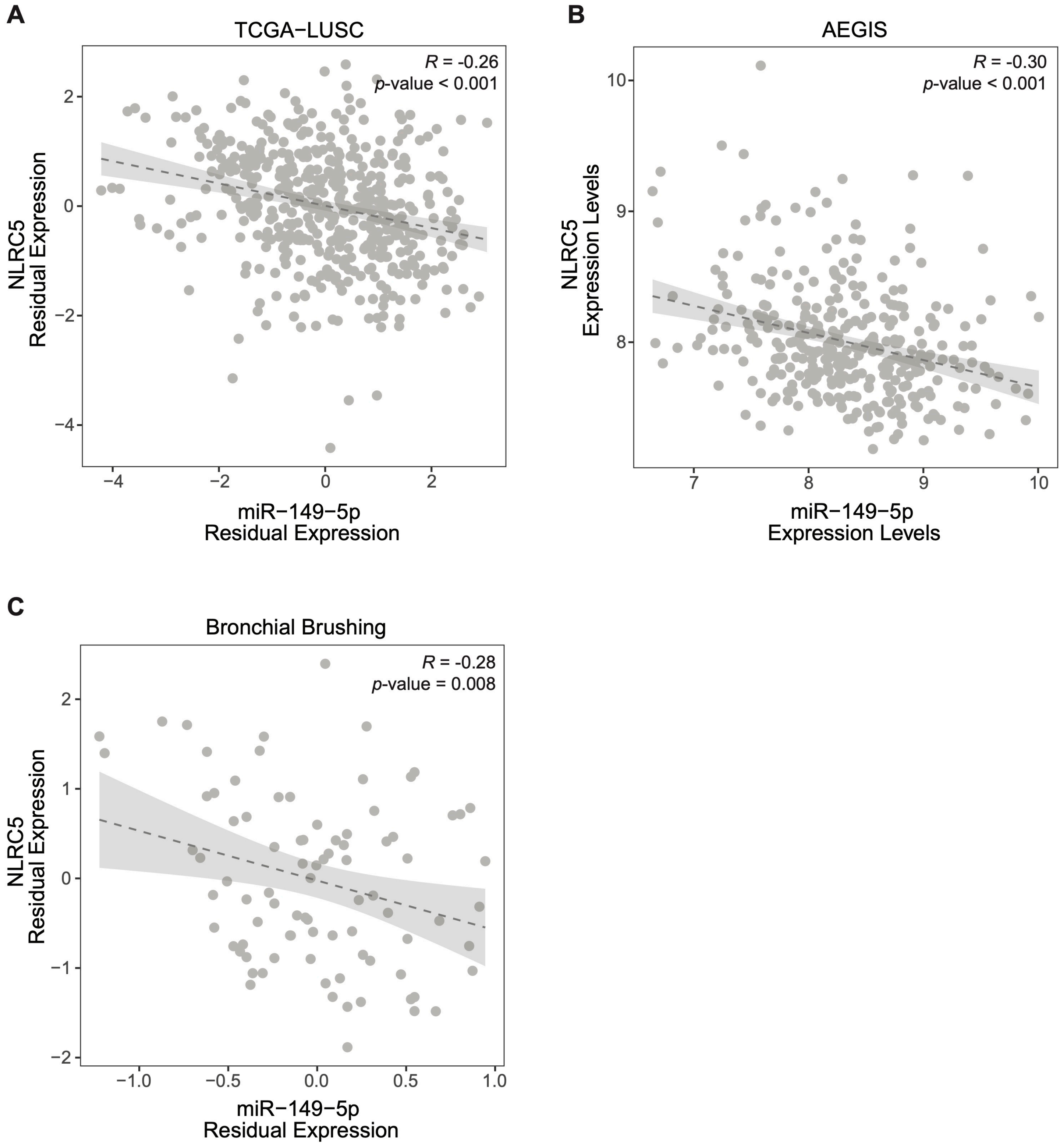


**Extended Data Figure 3. Expression level of hsa-miR-149-5p was significantly negatively correlated with that of NLRC5 in lung-related datasets.** Scatterplots showing the Pearson correlation between the expression levels of hsa-miR-149-5p and NLRC5 across three datasets: (**A**) TCGA-LUSC primary tumor samples (n=475), (**B**) AEGIS bronchial brushing samples (n=341), and (**C**) bronchial brushing samples (n=87) from this study. The dashed line represents the linear regression fit and the shaded gray region indicates the 95% confidence interval. There was a significant negative correlation, calculated using Pearson correlation, between hsa-miR-149-5p and NRLC5 in all datasets (**A-C**).

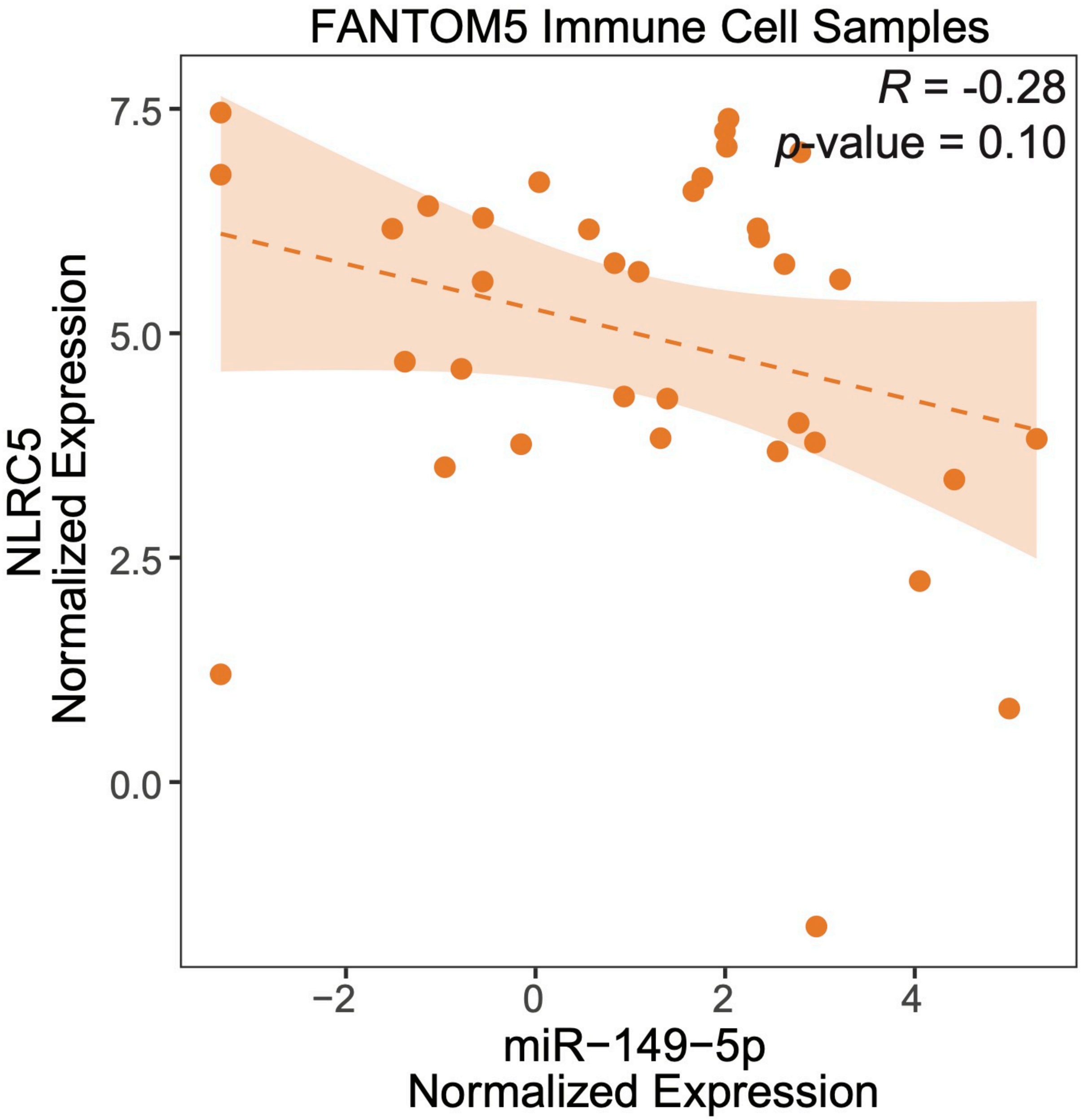


**Extended Data Figure 4. Correlation between hsa-miR-149-5p and NLRC5 expression levels within the samples from the FANTOM5 project.** Scatter plot of the Pearson correlation between the normalized expression levels of hsa-miR-149-5p and NLRC5 within the FANTOM5 samples derived from immune (n=37) cell compartments. The orange dashed line represents the linear regression fit and the shaded region indicates the 95% confidence interval.

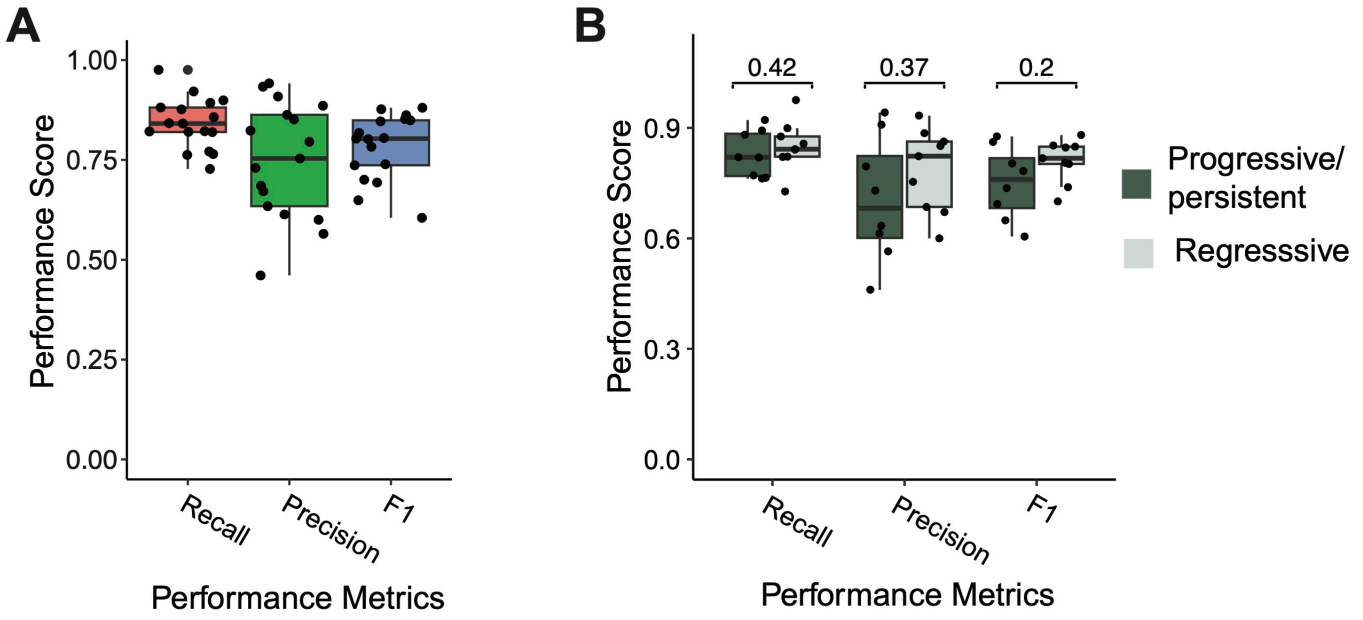


**Extended Data Figure 5. Performance of the hsa-miR-149-5p spot detection classifier.** (**A**) Boxplot showing the recall (red orange), precision (green), and F1(blue) scores for hsa-miR-149-5p detection classifier on regions in each miR-ISH image containing manual spot annotations (n = 19). (**B**) Boxplot showing the recall, precision and F1 scores for the hsa-miR-149-5p detection classifier between progressive/persistent PMLs (dark green, n= 10) and regressive PMLs (light green, n = 9). There are no significant differences in the performance metrics between progressive/persistent PMLs and regressive PMLs. Data indicate median with IQR, and whiskers indicate minimum and maximum measurement.

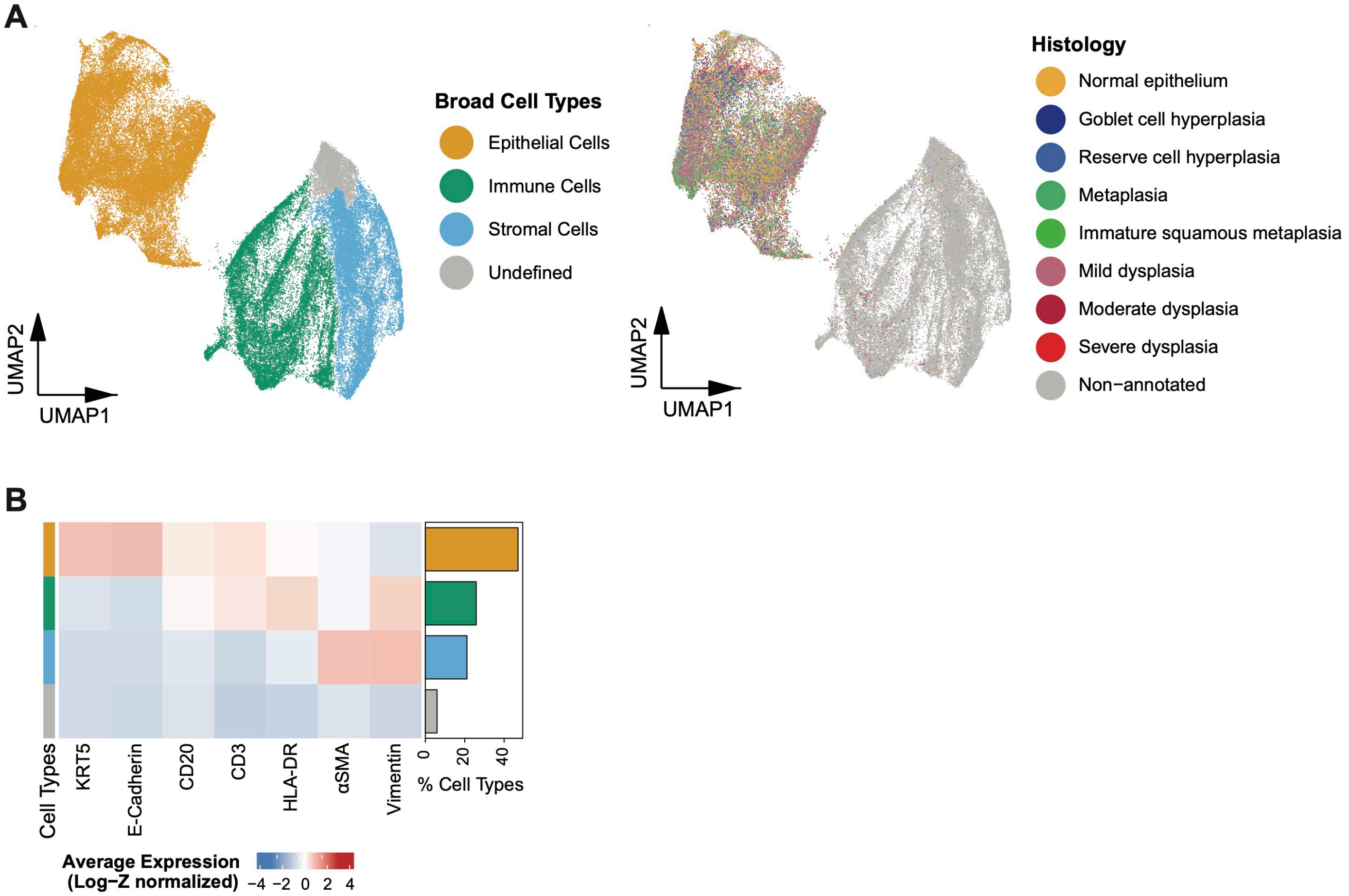


**Extended Data Figure 6. Identification of broad cell types in IMC data.** (**A**) Uniform Manifold Approximation and Projection (UMAP) plots showing 3 broad cell types (left) and histology annotations (middle) for all cells in the IMC images (n = 87,401). (**B**) Heatmap showing the average expression of canonical markers across the broad cell types with the bar plot showing the relative proportion of each cell type.

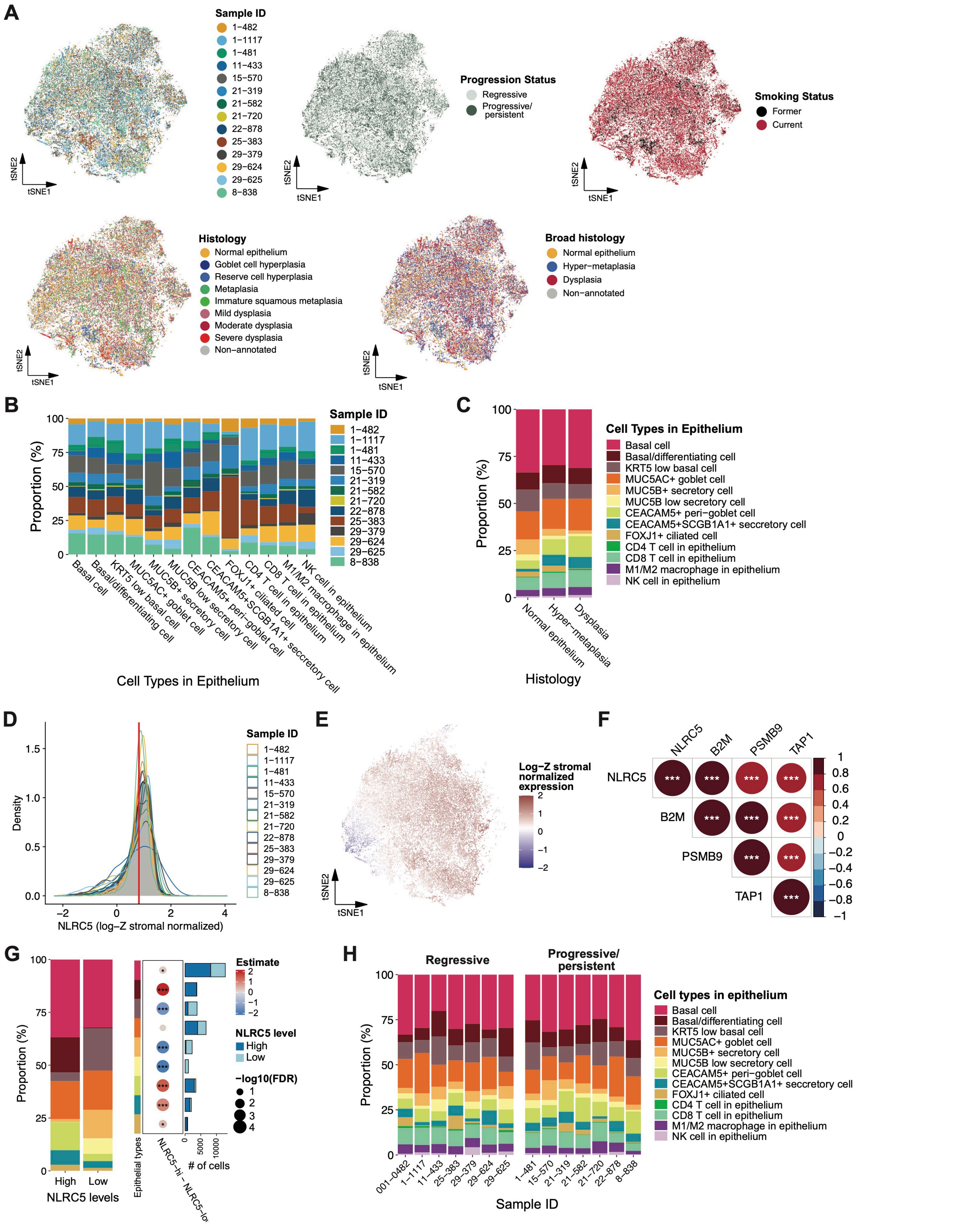


**Extended Data Figure 7. Analysis of IMC data from the epithelial tissue.** (**A**) tSNE visualization of cells within the epithelium (n = 41,147 cells) colored by sample ID (left top), progression status (middle top), smoking status (right top), histology (left bottom), and histology groups (middle bottom). (**B**) Stacked bar plot showing the relative proportion of cells contributed by each sample for each cell type identified within the epithelium. (**C**) Stacked bar plot showing the relative proportion of cell types across different histology groups in epithelium. (**D**) Density plots showing the distribution of the log-Z stromal normalized NLRC5 expression across epithelial cell populations for each sample. The gray shade shows the distribution of values across all samples and the red line represents the average of this distribution. The red line was used to dichotomize the cells within the epithelium into NLRC5-hi or NLRC5-low cells. (**E**) tSNE plot of cells within the epithelium colored by log-Z stromal normalized NLRC5 expression. (**F**) Bubble plot showing the correlation between NLRC5 expression and the expression of its downstream targets in epithelial cells. Color and dot size represent the Pearson correlation coefficient. (**G**) Stacked bar plot showing the relative proportions of epithelial cell types across NLRC5-high or NLRC5-low epithelial cells (left). Bubble plot showing the differential composition of 9 epithelial cell types between NLRC5-high and NLRC5-low epithelial cells. Bar plot to the right of the dot plot shows the number of cells in each cell type stratified by NLRC5-high versus low expression. Dot color represents the compositional estimate, and dot size represents the log FDR value computed by sccomp. (**H**) Stacked bar plot showing the relative proportion of cell types identified in epithelium for each sample stratified by PML outcome (regressive versus progressive/persistent). P values were FDR adjusted and determined by the t-test for correlation significance for (**F**), and the sccomp differential composition test for (**G**). * P <= 0.05, **P <= 0.01, ***P <= 0.001.

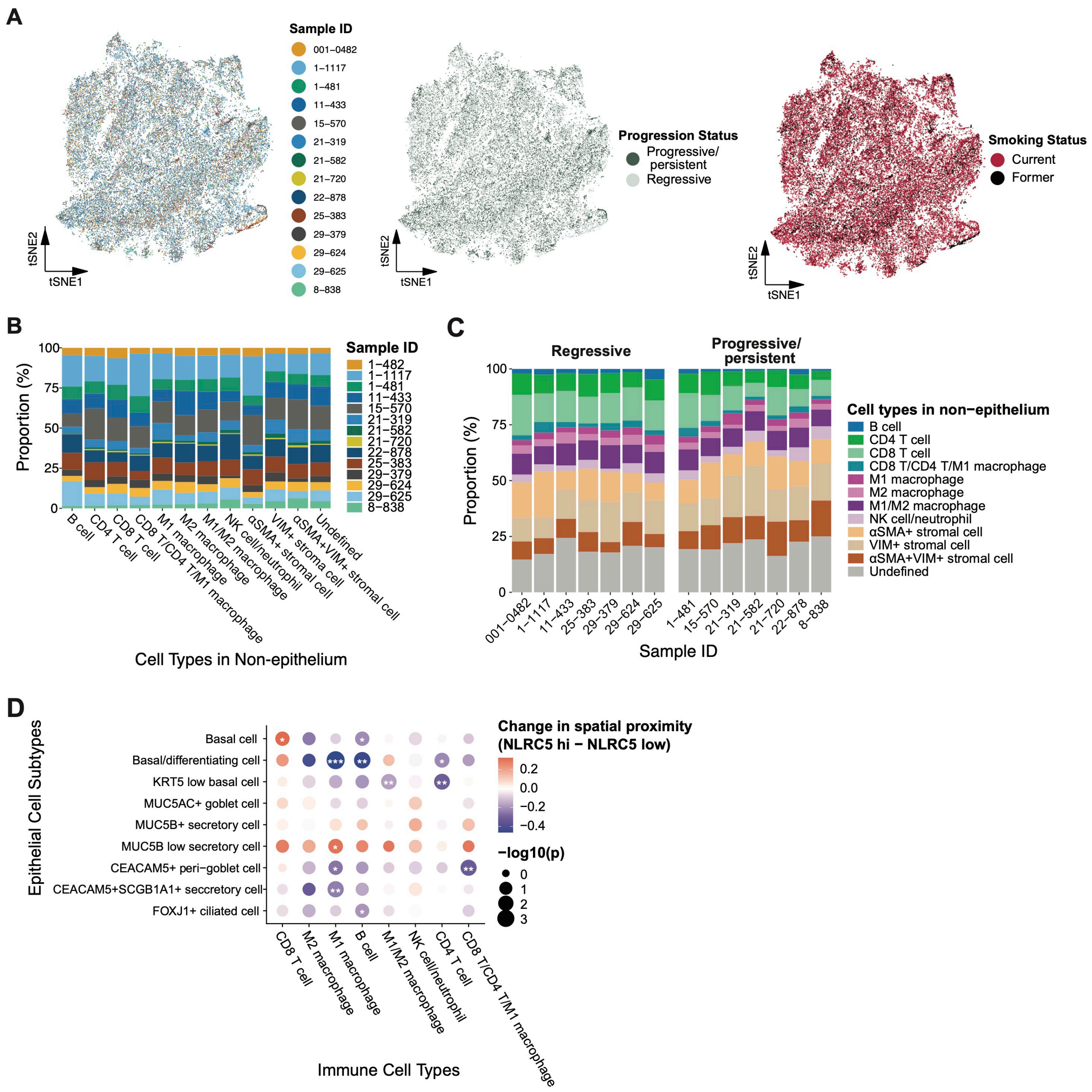


**Extended Data Figure 8. Analysis of IMC data from the non-epithelial tissue.** (**A**) tSNE visualization of cells not in epithelium (n = 46,254 cells) colored by sample ID (left), progression status (middle), and smoking status (right). (**B**) Stacked bar plot showing the relative proportion of cells contributed by each sample for each cell type identified within non-epithelium. (**C**) Stacked bar plot showing the relative proportion of cell types identified in the non-epithelium for each sample stratified by PML outcome (regressive versus progressive/persistent). (**D**) Bubble plot showing for each epithelial cell type the change in spatial proximity between NLRC5-high and NLRC5-low cells and immune cell populations. Dot color represents the changes in spatial proximity where red indicates closer spatial proximity between an immune cell population and NLRC5-high compared to NLRC5-low cells of a specific epithelial cell type, and dot size represents the log p-value. P values were determined by the two-sided paired Wilcoxon test for. * P <= 0.05, **P <= 0.01, ***P <= 0.001.

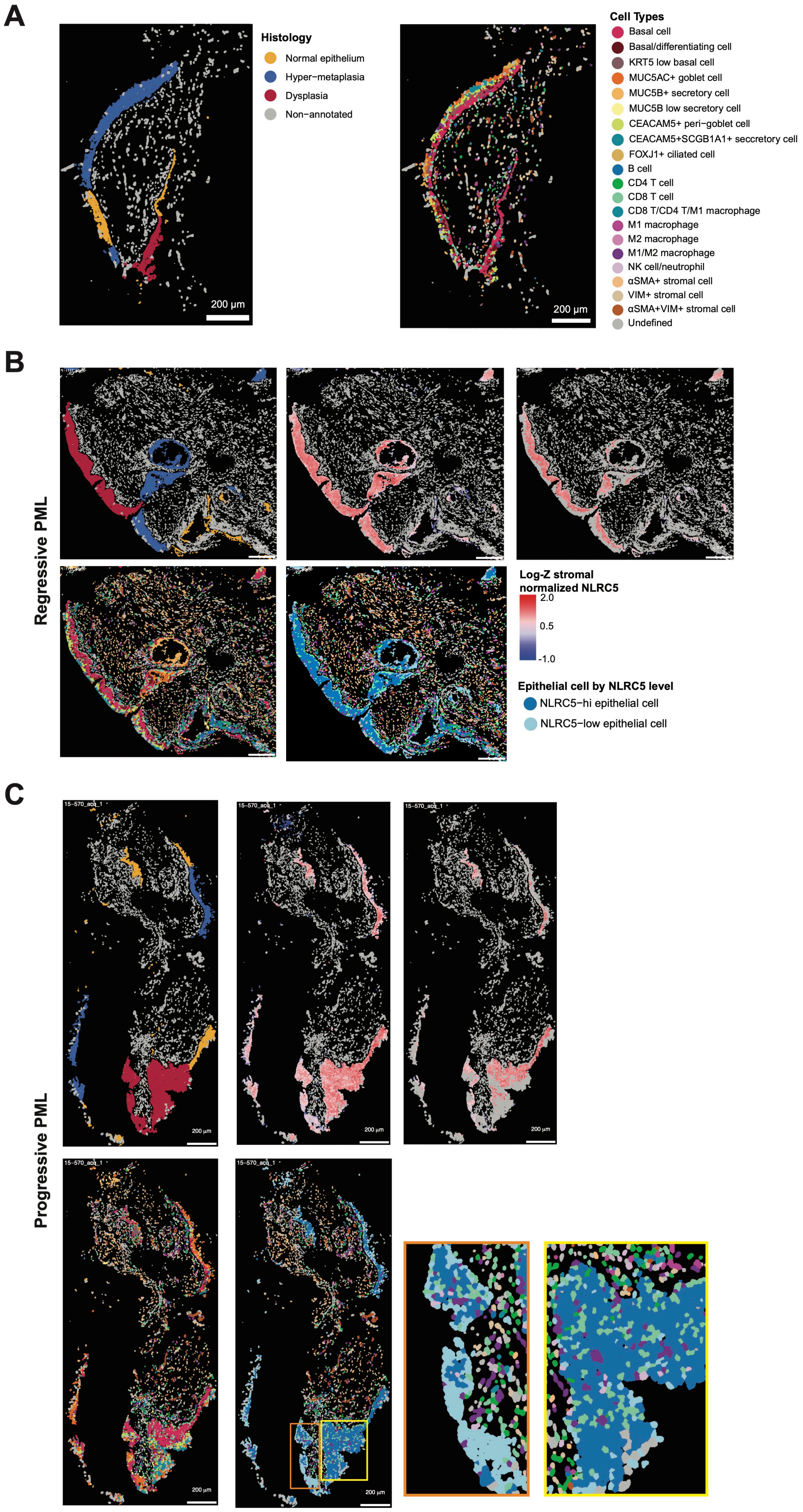


**Extended Data Figure 9. Representative IMC images.** (**A**) Representative IMC image where the segmented cells are shown colored by histology group (left) and cell type (right). (**B** and **C**)

Representative IMC images of a progressive PML (**B**) and a regressive PML (**C**) where segmented cells are colored by histology group (top left), NLRC5 expression of epithelial cells (top middle), NLRC5 expression of basal cells (top right), cell types (bottom left), and spatial proximity of NLRC5-high versus NLRC5-low cells with immune cells (bottom middle). In (**C**) the yellow and orange boxes in the bottom middle figure denote regions that are shown at a higher magnification.
